# Deep serum proteomics reveal biomarkers and causal candidates for type 2 diabetes

**DOI:** 10.1101/633297

**Authors:** Valborg Gudmundsdottir, Valur Emilsson, Thor Aspelund, Marjan Ilkov, Elias F Gudmundsson, Nuno R Zilhão, John R Lamb, Lori L Jennings, Vilmundur Gudnason

## Abstract

The prevalence of type 2 diabetes mellitus (T2DM) is expected to increase rapidly in the next decades, posing a major challenge to societies worldwide. The emerging era of precision medicine calls for the discovery of biomarkers of clinical value for prediction of disease onset, where causal biomarkers can furthermore provide actionable targets. Blood-based factors like serum proteins are in contact with every organ in the body to mediate global homeostasis and may thus directly regulate complex processes such as aging and the development of common chronic diseases. We applied a data-driven proteomics approach measuring serum levels of 4,137 proteins in 5,438 Icelanders to discover novel biomarkers for incident T2DM and describe the serum protein profile of prevalent T2DM. We identified 536 proteins associated with incident or prevalent T2DM. Through LASSO penalized logistic regression analysis combined with bootstrap resampling, a panel of 20 protein biomarkers that accurately predicted incident T2DM was identified with a significant incremental improvement over traditional risk factors. Finally, a Mendelian randomization analysis provided support for a causal role of 48 proteins in the development of T2DM, which could be of particular interest as novel therapeutic targets.

## Introduction

Type 2 diabetes mellitus (T2DM) is a progressive disease characterized by decreasing sensitivity of peripheral tissues to plasma insulin accompanied by compensatory hyperinsulinemia, and a gradual failure of the pancreatic islet β-cells to maintain glucose homeostasis. The worldwide prevalence of diabetes is projected to increase from 451 million in 2017 to 693 million by 2045^1^. In the past decade, the use of data-driven omics technologies has led to a significant advancement in the discovery of new biomarkers for complex disease. More than 240 genetic loci have been associated with T2DM^2–6^ and recent efforts utilizing genome-wide polygenic risk scores have shown a promising ability to predict those at risk of developing the disease^6,7^. Blood-based biomarker candidates with prognostic value for T2D have begun to emerge, such as the branched-chain amino acids (BCAAs) and other metabolites^8,9^. However, only fragmentary data are available for protein biomarkers for prediction of incident T2DM^10^. In fact, robust molecular biomarkers are yet to be established that add a clinically useful predictive value over glycemia markers such as fasting glucose and HbA1c^10^. Thus, identification of novel biomarkers for T2DM is crucial for early and improved risk assessment of the disease beyond what can be achieved through the use of conventional measures of glycemia and adiposity.

Proteins are the key functional units of biology and disease, however, high throughput detection and quantification of serum proteins in a large human population has been hampered by the limitations of available proteomic profiling technologies. The Slow-Off rate Modified Aptamer (SOMAmer) based technology has emerged as a powerful proteomic profiling platform in terms of sensitivity, dynamic range of detection and multiplex capacity^11–13^. A custom-designed SOMAscan platform was recently developed to measure 5,034 protein analytes in a single serum sample, of which 4,782 SOMAmers bind specifically to 4,137 distinct human proteins^14^. We applied this platform to 5,457 subjects of the Age, Gene/Environment Susceptibility (AGES)-Reykjavik study, a prospective study of deeply phenotyped subjects over 65 years of age^14,15^. In the present study we demonstrate the identification of novel serum protein biomarkers for incident and prevalent T2DM through logistic regression and LASSO penalized logistic regression analysis combined with bootstrap resampling. Finally, by applying a Mendelian Randomization (MR) analysis, we identify a subset of those proteins that may be causally related to T2DM.

## Results

The baseline characteristics of the population-based AGES-Reykjavik cohort participants with complete data for the current study (n = 5,438) are shown in **Table S1** and an overview of the cohort and study workflow is shown in **Fig. S1**. The full cohort with baseline measurements included 654 prevalent T2DM cases and 4,784 individuals free of T2DM. Out of 2,940 individuals without diabetes at baseline who participated in the 5-year AGESII follow-up visit (Methods), 112 developed T2DM within the period based on self-report, medication and/or fasting glucose measurement. As an internal validation cohort for incident T2DM, we considered the 1,844 AGES participants who were non-diabetic at baseline but did not participate in the AGESII 5-year follow-up visit, for whom we defined incident T2DM from prescription and medical records only (see Methods), resulting in 46 cases within up to a 12.8 years follow-period. As expected, both prevalent and incident T2DM cases differed markedly from individuals free of diabetes in terms of metabolic phenotypes at baseline (**Table S1**).

### Serum protein profile of prevalent T2DM

To first describe the serum protein profile associated with prevalent T2DM, we compared 654 prevalent T2DM cases to 4,784 non-diabetic individuals. Using a logistic regression adjusted for age and sex, we identified 520 unique proteins that were significantly associated with prevalent T2DM after Bonferroni correction for multiple hypothesis testing (P_adj_ < 0.05), with the strongest associations observed for ARFIP2, MXRA8 and CPM (**Fig. 1a, Table S2**). In a second model including adjustment for body mass index (BMI), 322 proteins remained statistically significant (**Table S2**). Many of the proteins were inter-correlated, with pairwise Pearson’s *r* ranging from −0.60 to 0.97 (**Fig. S2a**). A pathway and gene ontology (GO) enrichment analysis of all 520 proteins associated with prevalent T2DM revealed an enrichment of proteins involved in extracellular matrix (ECM)-receptor interaction, complement and coagulation cascades, metabolic processes and extracellular region (**Fig. S3a, Table S3**). We furthermore found the genes encoding the 520 prevalent T2DM-associated proteins to be enriched for high expression in liver, followed by other tissues that included kidney, gastrointestinal tract and pancreas (**Fig. S4a**). Thus, the diabetic state is reflected in a major shift in the serum proteome that is involved in metabolic, inflammatory and ECM processes.

**Fig. 1.**
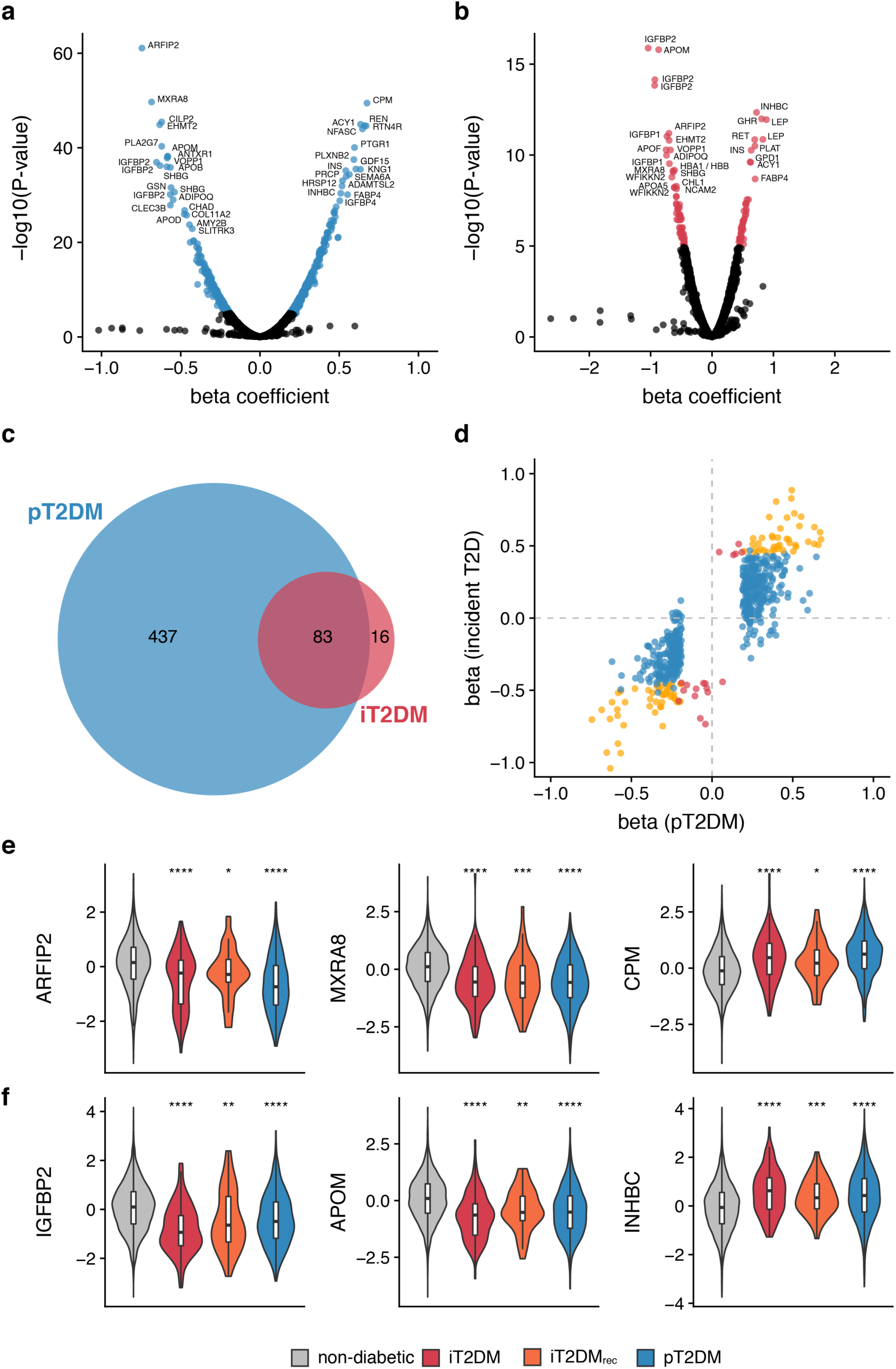
**a)** Volcano plots demonstrating serum protein (SOMAmer) associations with prevalent T2DM and **b)** incident T2DM. Points are colored where P_adj_ < 0.05. **c)** Venn diagram showing the overlap between unique proteins associated with prevalent T2DM (blue) and incident T2DM (red). **d**) Beta coefficients for associations between proteins (SOMAmers) and prevalent T2DM (x-axis) and incident T2DM (y-axis) The colors denote significant associations with prevalent T2DM (blue), incident T2DM (red) or both (yellow). **e)** Violin and boxplots showing serum protein levels across the AGES cohort stratified by T2DM status for top three proteins associated with prevalent T2DM and **f)** incident T2DM. Stars denote significant difference compared to the non-diabetic group with nominal P-values (two-sided t-test) as such: *P ≤ 0.05, **P ≤ 0.01, ***P ≤ 0.001, ****P ≤ 0.0001. Boxplots indicate median value, 25^th^ and 75^th^ percentile, whiskers extend to smallest/largest value no further than 1.5×IQR, outliers not shown. pT2DM, prevalent T2DM; iT2DM, incident T2DM in participants with AGESII follow-up visit; iT2DM_rec_, incident T2DM in participants without AGESII follow-up visit.

### Serum protein profile of incident T2DM

The serum protein profiles of T2DM patients observed in the cross-sectional analysis described above may represent shifts that occurred either before or after the onset of the disease. To identify serum protein signatures that preceded the onset of T2DM, we next focused our analysis on the 2,940 non-diabetic AGES participants who participated in a second study visit (AGESII) 5-years after the baseline visit, of which 112 developed T2DM within the follow-up period. In a logistic regression analysis adjusted for age and sex, we identified 99 unique proteins significantly associated with incident T2DM after Bonferroni correction for multiple hypothesis testing with the strongest associations observed for IGFBP2, APOM and INHBC (**Fig. 1b, Table S4**). After further adjustment for BMI, 24 proteins remained statistically significant (P_adj_ < 0.05) (**Table S4**). Once again we observed extensive correlations between many of the serum proteins, with pairwise Pearson’s *r* ranging from −0.55 to 0.97 (**Fig. S2b**). The majority (84/99 proteins or 85%) of proteins associated with incident T2DM were also associated with prevalent T2DM (**Fig. 1c**), an overlap that was highly significant (Fisher’s exact test P = 7.2×10^−63^), and the direction of effect was generally consistent (Spearman’s correlation coefficient = 0.82, **Fig. 1d-f**). The proteins associated with incident T2DM included proteins with an established role in T2DM (IGFBP2, adiponectin and insulin), proteins encoded by genes reported as T2DM GWAS loci^6^ (ATP1B2, PTPRS) and various apolipoproteins (APOM, APOF, APOA5). Functional enrichment analysis of the full set of 99 proteins associated with incident T2DM revealed a significant enrichment for numerous GO terms related to metabolism, lipid transport and response to insulin while enriched pathways included leptin signaling and adipogenesis (**Fig. S3b, Table S3**). Tissue expression enrichment analysis revealed a strong enrichment for genes expressed in liver, followed by adipose tissue (**Fig. S4b**). Thus, the functional annotation of the serum proteins associated with incident T2DM was characterized by tissue specific signatures and pathways that seem to reflect dyslipidemia and insulin resistance, which are critical in the development of T2DM. We compared our findings with previously described protein biomarker candidates for incident T2DM as previously reviewed^11^. Of 58 previously suggested candidates that were targeted in our study, we found 26 to be at least nominally associated (P < 0.05) with incident T2DM in our data and additional 15 with prevalent T2DM (**Table S5**).

### Predictive performance of protein biomarkers for incident T2DM

As it is of considerable interest to define a set of biomarkers for clinical prediction of T2DM, we aimed to define the subset of proteins associated with incident T2DM that had the best predictive value. To evaluate the power to discriminate between incident T2DM cases and non-cases, we applied a receiver operating characteristic (ROC) curve to compute the area under the curve (AUC). The AUC for incident T2DM using age and sex alone was 0.56 (95% CI 0.51-0.62) and a clinical model including the Framingham-Offspring Risk Score (FORS)^16^ components (age, sex, parental history of diabetes, BMI, systolic blood pressure, HDL, triglycerides, fasting glucose and abdominal circumference as a proxy for waist) yielded an AUC of 0.86 (95% CI 0.83-0.90). Only a single protein (REN) added significantly to the FORS model C-statistic (C_increase_ = 0.0055, P = 0.041, **Table S4**), thus motivating a multivariate predictor analysis. For this purpose, a least absolute shrinkage and selection operator (LASSO) logistic regression model combined with bootstrap resampling was fitted using incident T2DM as outcome and age, sex, and the full set of 4,782 SOMAmers as predictors. Here, a set of 32 non-zero parameter estimates gave the highest AUC when the tuning parameter log(lambda) was −4.54 for incident T2DM (**Fig. S5**). To account for randomness in the selection process, model performance and improved variable selection, the LASSO was bootstrapped 1,000 times through resampling. The proteins were rank-ordered with respect to how often they were selected during the bootstrap resampling and for the strength of association to incident T2DM in the logistic regression analysis. The top 20 protein predictors among those significantly associated (P_adj_ < 0.05) with incident T2DM in the logistic regression analysis are listed in **Table 1**. We investigated the added value of both top 10 and 20 ranked serum proteins beyond age, sex and the full FORS model. Both sets of proteins increased the predictive value significantly, where an addition of 10 and 20 proteins increased the AUC from 0.86 for the FORS model to 0.90 (P = 3.2×10^−3^) and 0.91 (P = 2.8×10^−4^), respectively (**Fig. 2a-b, Table 2**). The observed increase in AUC was considerably greater than for randomly sampled sets of proteins (**Fig. 2b**). Observed and predicted proportion with incident T2DM in each risk decile of the 20 protein discrimination model are shown in **Fig. S6**.

**Table 1.**
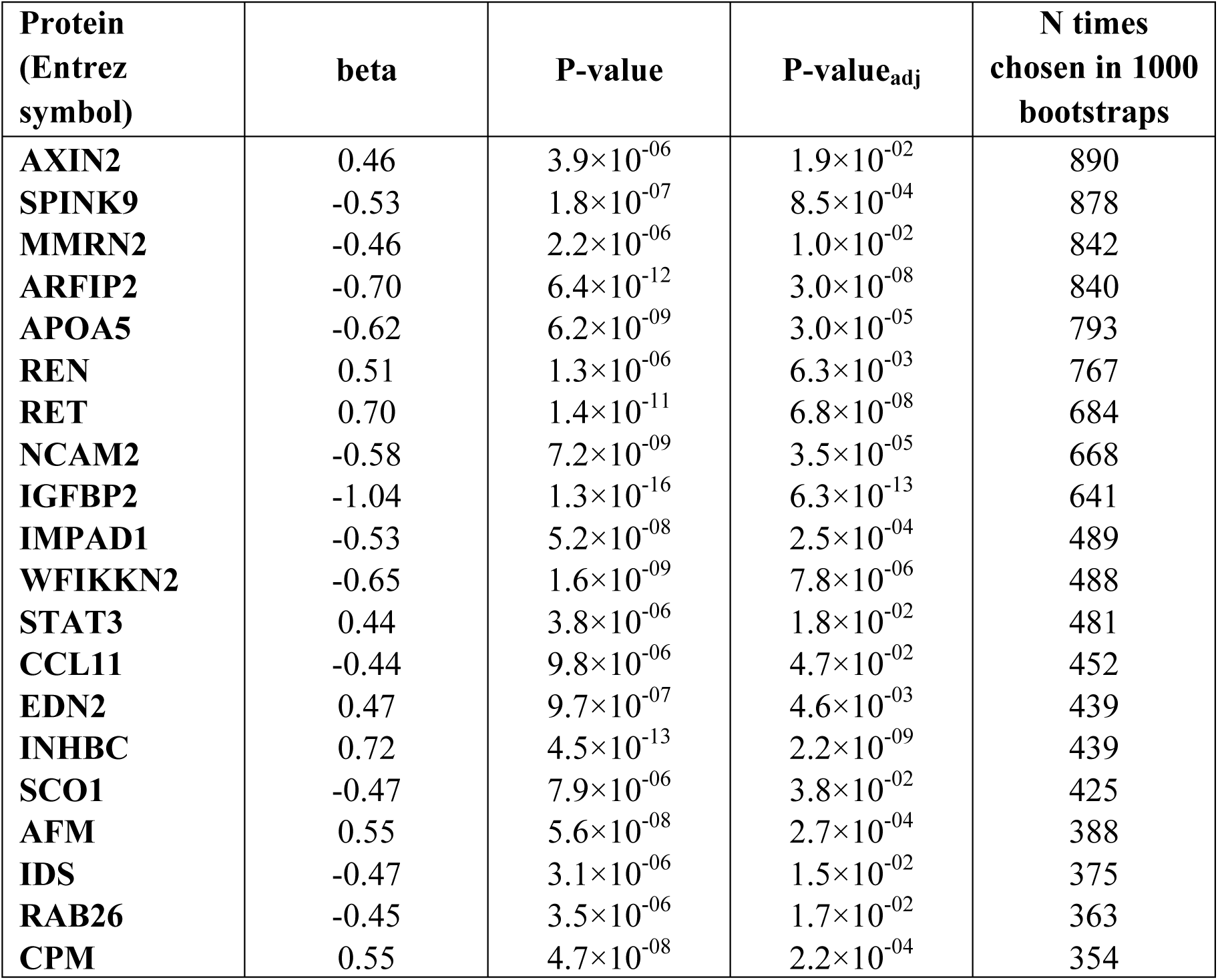
The top 20 proteins predicting incident T2DM as ranked by the number of times chosen in LASSO bootstrap analysis score in the AGES discovery sample (n = 2,940), shown with beta-coefficient, P-values and Bonferroni-corrected P-value from the logistic regression analysis adjusted for age and sex.

**Table 2.** Receiver operating characteristic discrimination scores (AUC) based on 10 or 20 top ranked protein predictors for incident T2DM together with baseline and the FORS clinical model in the AGES discovery sample (n = 2,940). P-values (paired two-sided DeLong’s test) are shown for the comparison of ROC curves to either the previous model or the baseline model.

| Model | N proteins | AUC | Lower bound | Upper bound | P-value previous | P-value baseline |
| --- | --- | --- | --- | --- | --- | --- |
| Age, sex | 0 | 0.56 | 0.50 | 0.61 | 1 | 1 |
| | 10 | 0.84 | 0.80 | 0.88 | $1.93 \times 10^{-20}$ | $1.93 \times 10^{-20}$ |
| | 20 | 0.87 | 0.83 | 0.90 | 0.016 | $2.63 \times 10^{-23}$ |
| FORS | 0 | 0.86 | 0.83 | 0.90 | 1 | 1 |
| | 10 | 0.89 | 0.86 | 0.93 | $3.25 \times 10^{-03}$ | $3.25 \times 10^{-03}$ |
| | 20 | 0.91 | 0.88 | 0.94 | 0.045 | $2.83 \times 10^{-04}$ |

**Fig. 2.**
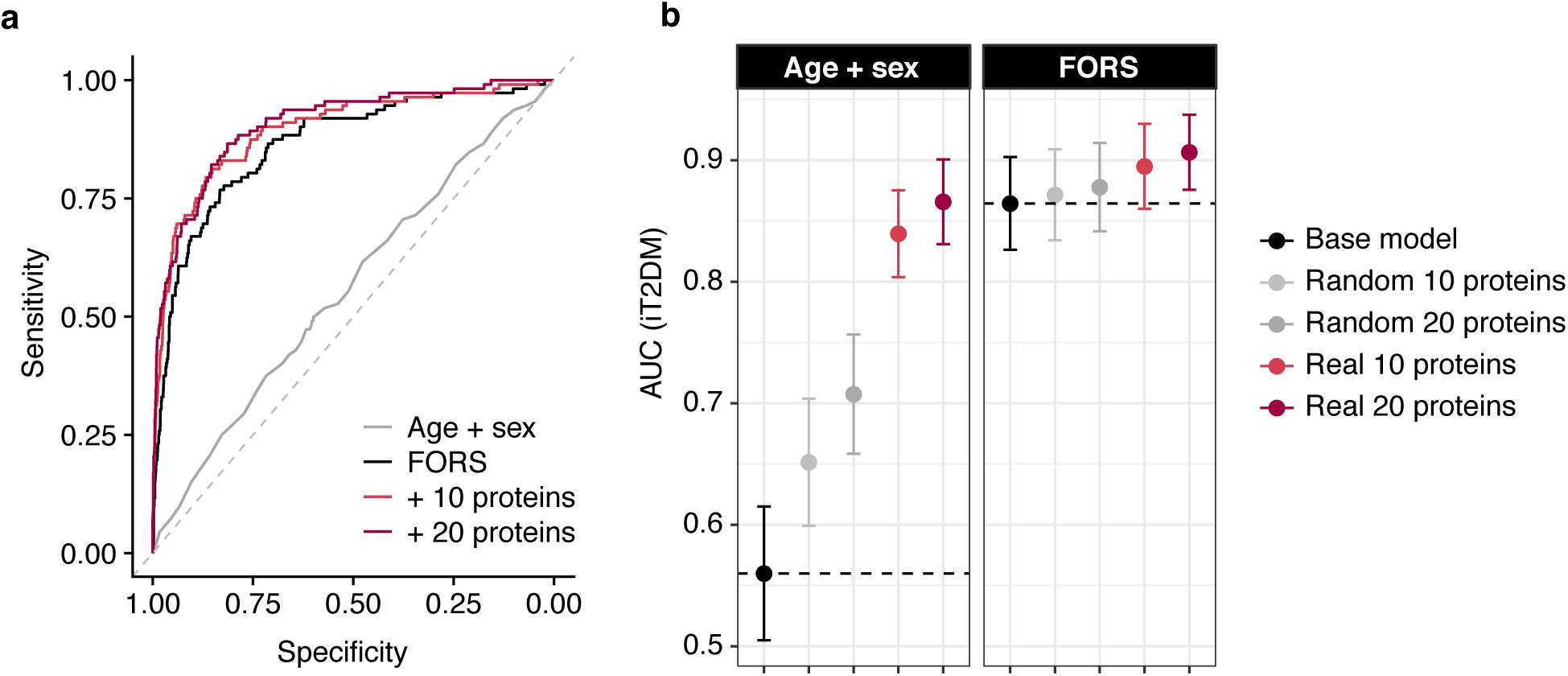
**a)** ROC curves showing the added value of top 10 and 20 ranked proteins (red shades) for prediction of incident T2DM compared to age and sex (grey) and Framingham-Offspring risk score, FORS (black) in the AGES cohort (n = 2,940, n_FORS_ = 2,926). **b)** The AUC for top 10 and 20 proteins (red shades) is shown compared to a base model (black point and dotted line) of age and sex (left) or FORS (right) and compared to the AUC obtained by 100 permutations of randomly sampled sets of proteins (grey shades). Error bars represent 95% confidence intervals.

To our knowledge, similar data does currently not exist in another cohort for independent replication of our findings. However, in addition to the bootstrap approach employed for internal validation, we performed a secondary validation approach using data from the 1,844 AGES-Reykjavik participants who were non-diabetic at baseline but did not participate in the AGESII 5-year follow-up visit and were thus not included in the discovery analysis for incident T2DM (**Table S1**). Using the 20 proteins chosen from the LASSO analysis (**Table 1**), the AUC for incident T2DM (as defined from prescription and medical records) was significantly increased from 0.80 for the FORS model to 0.84 (P = 6.6×10^−3^) (**Fig. S7, Table S6**) in this set of individuals.

### Potentially causal associations between protein biomarkers and T2DM

While it is not a requirement for clinically useful biomarkers to be causally related to disease, identifying causal disease pathways provides important insights for the development of new therapeutic strategies. We therefore performed a MR analysis^17^ to identify proteins with a potentially causal role in the development of T2DM (**Fig. S8**). To maximize the protein coverage for this analysis, we used a subset of the AGES cohort with available genetic data (n = 3,219) to select genetic instruments for the proteins of interest but note that *cis*-pQTLs identified in AGES replicated over 80% of *cis*-pQTLs reported by others^14^. For the genes encoding the 536 proteins associated with either incident or prevalent T2DM in our study, using a *cis*-window of 100 kb up-and downstream and including the exons and introns of the genes in question, we identified suitable genetic instruments (see Methods) for 246 proteins, of which 184 (75%) proteins had more than one independent (r^2^ < 0.1) instrument (**Table S7**). On average, we identified 5 (range 1 - 20) genetic instruments per protein (**Fig. S9**), which explained on average 6% (range 0.4% - 48%) of the variance in their respective protein levels and with a mean F-statistic of 85 (range 10 - 3014). Of note, the genetic variants regulating the levels of the T2DM-associated proteins were strongly enriched within enhancer regions mapped in liver and hepatocytes from the Encode and Roadmap consortia (**Fig. S4c-d**), supporting the previously observed enrichment for liver expression of the genes encoding the T2DM-associated proteins.

We performed a two-sample MR analysis, integrating the genetic instruments for protein levels identified in AGES with summary statistics from the recent DIAMANTE GWAS for T2DM in 898,130 European individuals (74,124 T2DM cases and 824,006 controls)^6^. In this analysis, 48 proteins were supported as potentially causal (P < 0.05) for T2DM with the strongest support for MMP12, HIBCH and WFIKKN2 (**Fig. 3, Fig. S10**). Of these 48 proteins, few exhibited evidence of heterogeneity (2/36 proteins with >1 instrument) or pleiotropy (1/30 proteins with >2 instruments) (**Table S8)**. Proteins for which multiple genetic instruments were available tended to have smaller estimated effect sizes, together with a narrower confidence interval (**Fig. 3**). Of the 48 proteins, three (WFIKKN2, INHBC and AFM) were among the 20 proteins selected for the prediction of incident T2DM in the LASSO analysis. We further tested the 48 potentially causal proteins in a one-sample MR analysis using data from 3,196 AGES participants with available genotype data (N_T2DM_ = 368, 11.5%), fitting an age and sex adjusted two-stage regression with the second stage as a logistic regression. Using this approach, we obtained additional support (P < 0.05 and directionally consistent estimates) for four proteins (RBP7, IL18R1, FAM177A1, AFM) (**Fig. S11, Table S8**). We compared the observational and MR estimates for all 48 proteins (**Table S8, Fig. S12**). As expected due to a small sample size, the one-sample MR estimates were less precise than the two-sample MR estimates, as illustrated by the wider confidence intervals. We observed directional consistency between observational and two-sample MR estimates for 26 out of 48 (54%) proteins (**Table S8**), which neither related to the strength of the MR associations nor the number of instruments per protein (**Fig. S13**). As an example of discrepancies between observational and MR estimates, we found serum levels of MMP12 to be increased in T2DM, consistent with previous reports^18^, whereas the MR estimate for MMP12 suggested a protective effect for T2DM. These findings are similar to the reported protective MR estimate for MMP12 and risk of coronary heart disease^19^ whereas clinical and experimental studies have shown higher levels of MMP12 in cardiovascular disease^18,20^.

**Fig. 3.**
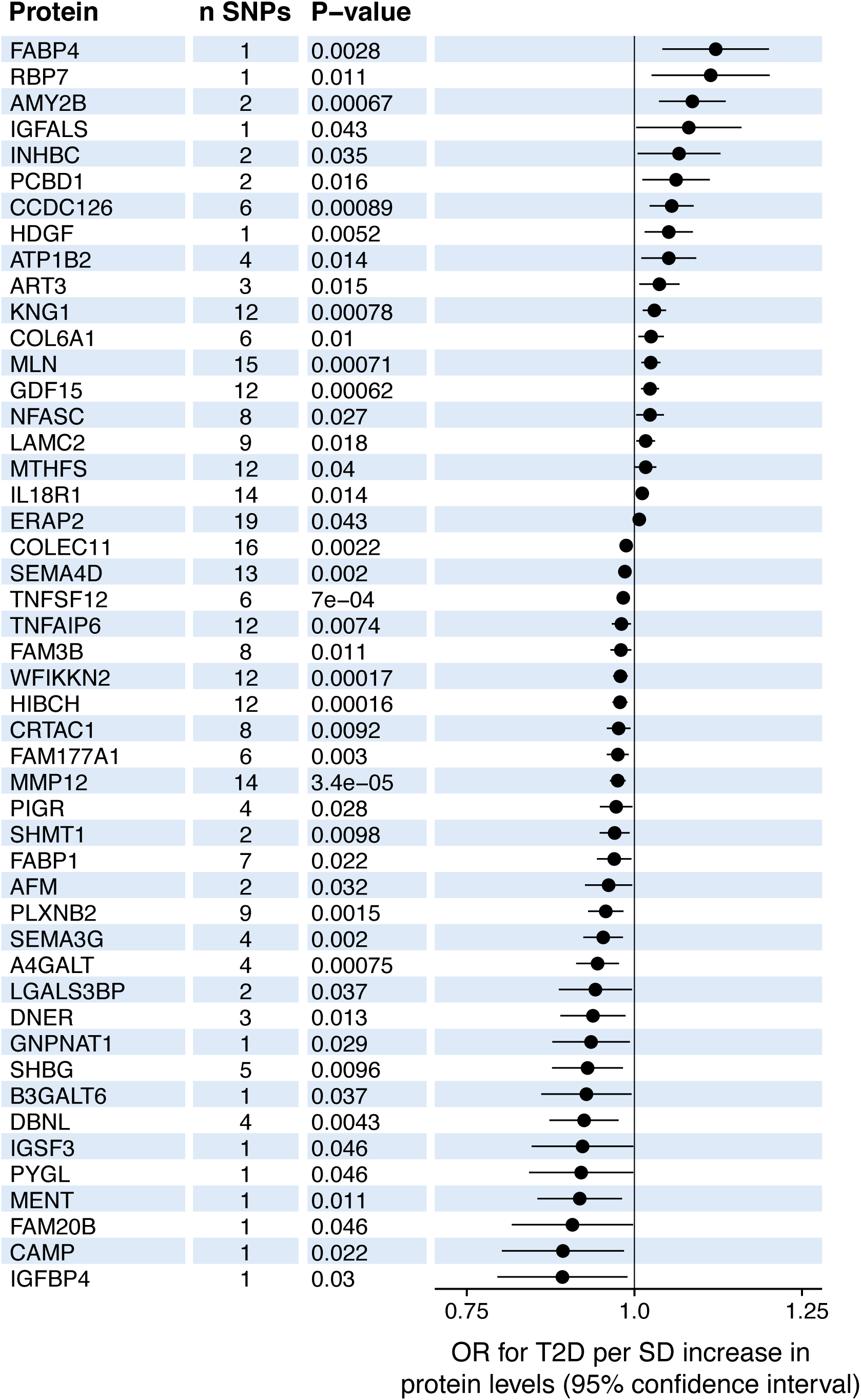
Forest plot for the 48 proteins supported as causal (P < 0.05) in the two-sample MR analysis, together with the number of SNPs used as instruments and the MR P-value. MR estimates were obtained using the inverse variance weighted method when >1 SNP was available for a given protein, but otherwise with the Wald ratio.

## Discussion

To our knowledge, the primary data used in the present study is the largest protein dataset generated to date in terms of number of proteins measured and human samples screened. In the literature there are few descriptions of plasma protein based biomarkers and drug targets for incident T2DM, and those available are usually limited to relatively few protein measurements^21–25^. In this study of a population-based sample of 5,438 elderly Icelanders, we advance the current knowledge by describing hundreds of proteins significantly associated with prevalent or incident T2DM, or both.

The large number of proteins significantly associated with prevalent T2DM demonstrates a major shift in the serum proteome in the diabetic state. We note that we have previously shown that the time between diagnosis and sample collection had no effect on the association of individual proteins to prevalent disease^14^. This proteomic shift seems to some extent be driven by inflammatory processes and ECM alterations given the observed enriched pathways. By contrast, these pathways were not enriched among proteins associated with incident T2DM, suggesting they may be secondary to the onset of the disease. Further studies of these proteomic changes are required to understand if and how they may affect downstream complications of T2DM, as diabetes-induced changes of the ECM may for example contribute to cardiovascular disease^26^. While we observed some proteomic changes specific to prevalent T2DM, others could be observed before the onset of the disease as illustrated by a large subset of the proteins also being associated with 5-year incident T2DM in non-diabetic individuals. A BMI-adjusted model suggested that a considerable proportion of the proteins were associated with T2DM via obesity. The proteins associated with incident T2DM were mainly involved in lipid transport, metabolism and insulin response, supporting the involvement of these pathways during the preclinical stage of T2DM. Both sets of proteins associated with prevalent or incident T2DM were enriched for liver-specific gene expression compared to the full set of 4,137 serum proteins measured, consistent with the genetic variants regulating their levels being enriched in enhancers mapped in liver tissue and hepatocyte cell lines. These results underscore that the diabetic serum proteomic signatures seem to reflect processes ongoing in the liver, although other tissues also contribute to the proteomic changes related to T2DM, as demonstrated for example by the enrichment of adipose expression among proteins associated with incident T2DM.

A systematic review of blood-borne and urinary biomarkers for incident T2DM concluded that no single marker has been identified with a prediction value comparable to that of glycemia markers, although some can add value to the prediction^10^, thus highlighting a potential need for multivariate predictors. A major strength of our study is the extensive protein coverage of the applied array, making this the most comprehensive screening of serum proteins for prediction of incident T2DM to date. Through LASSO regression we identified a subset of 20 proteins that as a group added significantly to the FORS model of clinical variables for prediction of incident T2DM, both in an internal bootstrap validation setup and importantly also in a separate sample of the AGES cohort that was not used for discovery analysis. However, it should be noted that our validation sample contained few cases and different criteria were applied to define incident cases than for the discovery sample, since the validation sample did not include a fasting glucose measurement and thus did not capture undiagnosed or non-medicated individuals. It may however also be considered a strength that the protein predictors still improved the prediction of incident T2DM despite these differences but we acknowledge that these efforts serve as internal validation only. Currently, similar data in other cohorts are lacking and future efforts will have to be made for replication of our findings in independent populations and across different proteomics technologies.

The MR analysis revealed a total of 48 proteins that may be causally related to T2DM. Among the candidate proteins with the strongest support, we found HIBCH that is a BCAA catabolic enzyme, where the MR estimate suggested an inverse causal effect between the proteins and risk of T2DM. Circulating BCAAs levels have consistently been shown to predict T2DM^27^ although the underlying mechanisms are complex and remain to be fully understood^28^. Our findings support a model where higher protein expression of the BCAA catabolic pathway reduces risk of T2DM. Members of the PPAR signaling pathway (FABP4, FABP1) were also found among the causal candidate proteins for T2DM. PPARs are the target of the thiazolidinediones anti-diabetic drug class and our results suggest that other members of this pathway could be considered as therapeutic targets. In fact, FABP4 inhibitors have been proposed as novel therapeutic strategies for obesity and T2DM^29^ and a PPARg-regulated^30^ retinol-binding protein, RBP4, is similarly being considered as an anti-diabetic target^31^. Our results from both two- and one-sample MR analysis implicate another retinol-binding protein, RBP7, the expression of which is affected by PPARg ligands^32^, which may be an interesting novel candidate for follow-up studies.

Three proteins from the 20 protein predictor for incident T2DM were also supported as causal by the MR analysis; afamin (AFM), inhibin β_C_ (INHBC) and WFIKKN2. Afamin has been associated with both prevalent and incident T2DM in a large-scale pooled study of eight prospective cohorts^33^ and we here obtain support from both two-and one-sample MR analyses for it to play a causal role in the disease. Less is known about the function of the other two proteins; inhibin β_C_ is one of the inhibin/activin hormones and is highly expressed in liver whereas WFIKKN2 is known to bind GDF8/11 proteins with high affinity^34^, both of which have been implicated in diabetes^35,36^. We and others have shown that genetic variants in the *WFIKKN2* region regulate serum GDF8/11 levels in *trans* via WFIKKN2 protein levels^14,19^ and previously noted a correlation between WFIKKN2 and GDF8/11 serum levels^14^, however in the current study we did not find a significant association between GDF8/11 and T2DM so additional studies are required to understand the mechanisms by which WFIKKN2 may affect risk of T2DM.

The availability of both exposure and outcome data in our dataset provided the opportunity to compare causal estimates from the two-sample MR analysis and the observational estimate. In many cases we found these estimates to disagree. Inconsistent directionality between causal and observational estimates has been noted for particular serum proteins, such as for MMP12 and the risk of coronary heart disease^19^, for which we find a similar inconsistency with regard to T2DM. Further work will be required to understand the underlying causes of these inconsistent estimates, which indicate a complex relationship between genetics, protein mediators and disease.

To conclude, our results demonstrate a major shift in the serum proteome before and during the diabetic stage. The many signals observed in our study suggest that there is potential for developing clinically useful serum protein panels for T2DM risk prediction that can add information over traditional risk factors, thus promoting early diagnosis and improved prognosis of those at risk of developing the disease. Furthermore, proteins supported as potentially causal in our data could be of particular interest as novel therapeutic targets.

## Methods

### Study population

Cohort participants aged 66 through 96 were included from the AGES – Reykjavik Study^15^, a single-center prospective population-based study of deeply phenotyped subjects (n = 5,457). After excluding individuals without a fasting glucose measurement or with established type 1 diabetes, 5,438 individuals remained for analysis in the current study (mean age 76.6 ± 5.6 years). All AGES study cohort members were European Caucasians. Blood samples were collected at the AGES baseline visit after an overnight fast, serum was prepared using a standardized protocol and stored in 0.5 ml aliquots at −80°C. T2DM was determined from self-reported diabetes, diabetes medication use or fasting plasma glucose ≥ 7 mmol/L according to the American Diabetes Association guidelines^37^. Of the 4,784 AGES participants free of T2DM at first visit in AGES, 2,940 attended a 5-year follow-up visit (AGESII). Those with manifest T2DM at the five years follow-up visit were classified as incident T2DM cases, using same criteria as for the baseline visit. For the remaining 1,844 individuals who did not attend the AGESII follow-up visit, we used linked medical and prescription records and defined incident T2DM as having a registered ICD10 code starting with ‘E11’ or an ATC prescription code starting with ‘A10’ at any given time after the AGES baseline visit. Prescription records were obtained from a centralized database of drug prescriptions from the Directorate of Health in Iceland. Lipids, fasting glucose and HbA1c levels were measured on a Roche Hitachi 912 instrument, with reagents from Roche Diagnostics. Fasting insulin levels were measured on a Roche Elecsys 2010 instrument with an electrochemiluminescence immunoassay, using two monoclonal antibodies and a sandwich principle. The first IRP WHO Reference Standard 66/304 (NIBSC) was used to standardize the method. BMI was calculated as weight/(height)^2^. Abdominal circumference was measured in cm and used as a proxy for waist circumference in the FORS^16^ clinical model, as waist circumference was not measured at the AGES baseline visit. Parental history of diabetes was obtained from questionnaires administered at the baseline AGES visit.

The AGES-Reykjavik study was approved by the National Bioethics Committee in Iceland (approval number VSN-00-063), the National Institute on Aging Intramural Institutional Review Board (US), and the Data Protection Authority in Iceland. Informed consent was obtained from all study participants.

### Protein profiling platform

Each protein has its own detection reagent selected from chemically modified DNA libraries, referred to as Slow Off-rate Modified Aptamers (SOMAmers)^38^. We designed an expanded custom version of the SOMApanel platform to include proteins known or predicted to be found in the extracellular milieu, including the predicted extracellular domains of single- and certain multi-pass transmembrane proteins as previously described^14^. The new aptamer-based platform measures 5,034 protein analytes in a single serum sample, of which 4,782 SOMAmers bind specifically to 4,137 human proteins (some proteins are detected by more than one SOMAmer) and 250 SOMAmers that recognize non-human targets (47 non-human vertebrate proteins and 203 targeting human pathogens). Serum levels of 4,137 human proteins were determined at SomaLogic Inc. (Boulder, US) in distinct samples from 5,457 individuals essentially as previously described^12,14^. We note that albumin-tolerance testing is a part of standard assay development at SomaLogic and has been evaluated for all analytes on the new custom-designed aptamer-based platform, showing no effect of albumin addition on the SOMAmer-protein interactions. To avoid batch or time of processing biases, both sample collection and sample processing for protein measurements were randomized and all samples run as a single set. The 5,034 SOMAmers that passed quality control had median intra-assay and inter-assay coefficient of variation, 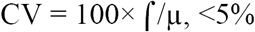, or similar to that reported on variability in the SOMAscan assays^38^. Finally, in addition to multiple types of inferential support for SOMAmer specificity towards target proteins including cross-platform validation and detection of the many *cis*-acting effects^14^, a direct measures of the SOMAmer specificity for 779 of the SOMAmers in complex biological samples was performed using tandem mass spectrometry^14^. Hybridization controls were used to correct for systematic variability in detection and calibrator samples of three dilution sets (40%, 1% and 0.005%) were included so that the degree of fluorescence was a quantitative reflection of protein concentration.

### Genotyping and imputation

For the MR analysis, we included 3,219 AGES participants for whom genetic data was available. Genotyping was performed using the Illumina 370CNV BeadChip array and genotype calling was performed using the Illumina Bead Studio. Samples were excluded based on sample failure, genotype mismatch with reference panel and sex mismatch on genotypes^39^. Imputation (1000 Genomes Phase 3 v5 reference panel) was performed using MaCH (version 1.0.16), and the following QC filtering was applied at the variant level: call rate (<97%), Hardy Weinberg Equilibrium (p < 1×10^−6^, PLINK mishap haplotype-based test for non-random missing genotype data (p < 1×10^−9^), and mismatched positions between Illumina, dbSNP and/or HapMap.

### Statistical analysis

Prior to the analysis of the protein measurements, we applied a Yeo-Johnson transformation on the protein data to improve normality, symmetry and to maintain all protein variables on a similar scale^40 35^. Logistic regression was run for all 4,782 SOMAmers targeting 4,137 human proteins for incident or prevalent T2DM as outcome, with age and sex included as covariates, and an additional model including BMI. Associations with P-value below a Bonferroni corrected threshold (P < 0.05/4,782 = 1.1×10^−5^) were considered significant. When more than one SOMAmer was available for the same protein, the one with the lowest P-value in the age and sex adjusted model was retained for all downstream analyses. Functional enrichment analysis for selected sets of proteins were performed using g:Profiler^41^, using the full set of human proteins targeted by the SOMApanel as background and a significance threshold of Benjamini-Hochberg FDR < 0.05. Tissue-specific gene expression enrichment analysis was performed using the TissueEnrich R package^42^.

For establishing a multivariate protein predictor for incident T2DM, we ran a Least Absolute Shrinkage and Selection Operator (LASSO) (L1-regularized regression) logistic regression model, with incident T2DM as outcome and age, sex, and proteins as predictors, using the glmnet R package for LASSO regression^43^. The LASSO solution is found by maximizing the diagnostic capacity of the predictors (the area under the curve or AUC) with constraints on the parameter estimates. With the LASSO approach most of the regression parameter estimates are set to zero. The constraint is chosen via cross-validation which introduces some randomness into the solution process. To account for the randomness in the selection process and to reduce chance of overfitting, the whole process was bootstrapped 1,000 times. The proteins selected for the final 10 and 20 protein predictors were chosen from those significantly associated with incident T2DM in the original logistic regression analysis, but ranked by the number of times they were chosen in the LASSO bootstrap analysis.

We assessed discrimination or differentiation between T2DM cases and non-cases through the receiver operating characteristic (ROC) curve, which is a graph of the true positive rate versus the false positive rate for each classification rule derived from a prediction model^44^. To quantify the predictive value of the selected set of proteins, the area under the ROC curve (AUC) was estimated. The AUC can be interpreted as the probability that a patient with the outcome is given a higher probability of the outcome by the model than a randomly chosen patient without the outcome^44^. ROC curves were compared with a paired two-sided DeLong’s test for two correlated ROC curves using the pROC package in R^45^.

For the MR analysis we identified genetic instruments as follows. For each protein, SNPs within a *cis* window of 100 kb up- or downstream of the respective protein-encoding gene (and including the gene in question) were tested for an association with protein levels in a linear regression model adjusted for age and sex assuming an additive genetic model. SNPs were included as genetic instruments if the association with protein levels was window-wide significant (P < 0.05/number of SNPs in the given window, similar to what was previously described^14^) and the F-statistic ≥ 10. Finally, the genetic instruments per protein were filtered to only include independent signals (r^2^ > 0.1, > 500 kb apart), identified using the clump_data command in the TwoSampleMR R package^46^ where linkage disequilibrium is calculated between the provided SNPs using European samples from the 1000 Genomes project and only the SNP with the lowest P-value retained among those in LD. We investigated cell-type specific enhancer enrichment of the genetic instruments compared to established GWAS loci through HaploReg v4.1^47^ using the SNP with the lowest association P-value per protein.

The two-sample MR analysis was performed using the “TwoSampleMR” R package^46^, using DIAMANTE GWAS summary statistics for T2DM without adjustment for BMI in European individuals^6^ as outcome. The inverse variance weighted method was used for the MR analysis unless only one genetic instrument was available, in which case the Wald ratio was used. A Cochran’s Q test (‘mr_heterogeneity’ function in the TwoSampleMR package) was used to evaluate heterogeneity of instruments and MR Egger regression (‘mr_pleiotropy_test’ function in the TwoSampleMR package) performed for indication of horizontal pleiotropy. For the one-sample MR analysis we performed a two-stage instrumental variable regression, with the second stage as a logistic regression, where a weighted genetic risk score was used as an instrumental variable when more than one genetic instrument was available for a given protein.

## Data availability

The custom-design Novartis SOMAscan is available through a collaboration agreement with the Novartis Institutes for BioMedical Research. Data from the AGES Reykjavik study are available through collaboration under a data usage agreement with the IHA. All data supporting the conclusions of the paper are presented in the main text and supplementary materials.

## Supplementary Table legends

**Table S1**. AGES-Reykjavik cohort baseline characteristics stratified by follow-up data availability and T2DM status.

**Table S2**. Serum protein associations with prevalent T2DM in the AGES cohort (n = 5,438). Results are shown for logistic regression models adjusted for age and sex, or age, sex and BMI. P_adj_, Bonferroni corrected P-value.

**Table S3**. Functional enrichment results from gProfiler for proteins associated with prevalent T2DM, incident T2DM or significant in the two-sample MR analysis for T2DM, using the full SOMApanel as background. Benjamini-Hochberg adjusted P-values <0.05 are highlighted in yellow.

**Table S4**. Serum protein associations with incident T2DM in the AGES cohort (n = 2,940). Results are shown for logistic regression models adjusted for age and sex; age, sex and BMI or the Framingham Offspring Risk Study (FORS) clinical model for prediction of incident T2DM. For the FORS model we show the AUC for the full model, together with the AUC increase for the given protein over the FORS clinical model alone and the respective P-value comparing the two (paired two-sided DeLong’s test).

**Table S5**. Overview of 57 published biomarker candidates for incident T2DM, together with the observed significance level of the corresponding protein measured in the current study. NS, not significant, P_adj_, Bonferroni adjusted P-value.

**Table S6**. Receiver operating characteristic discrimination scores (AUC) based on 10 or 20 top ranked protein predictors for incident T2DM together with baseline and the FORS clinical risk model in the AGES validation sample (n = 1,844). P-values (paired two-sided DeLong’s test) are shown for the comparison of ROC curves to either the previous model or the baseline model.

**Table S7**. An overview of the associations between the *cis*-SNPs used as instruments for the 246 proteins that were included in the MR analysis.

**Table S8**. Two- and one-sample Mendelian randomization results for 48 T2DM-associated proteins that were significantly (P < 0.05) associated with T2DM in the two-sample MR analysis.

**Fig. S1.**
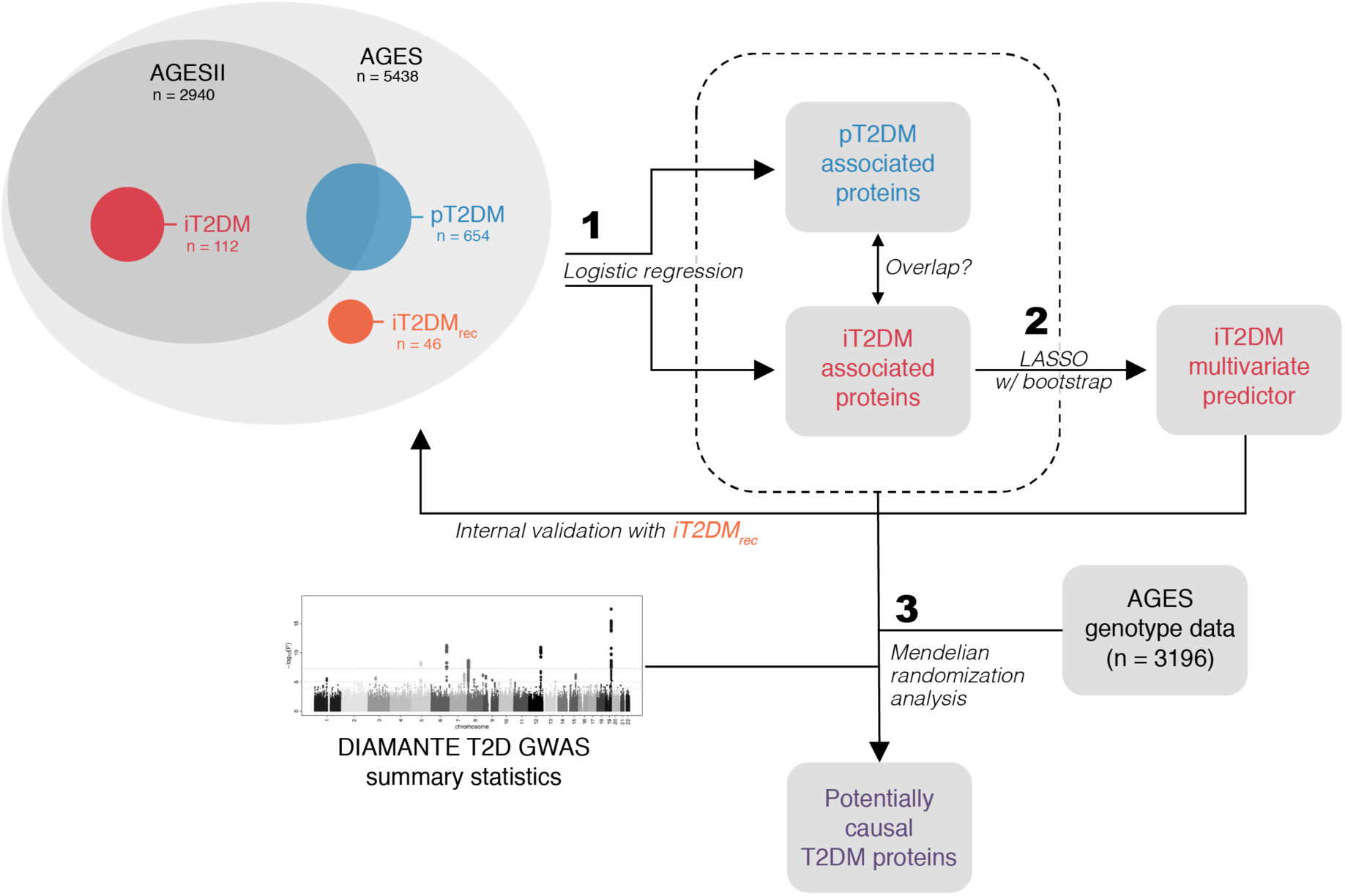
Workflow of the current study. The top left Venn diagram provides an overview of the AGES cohort, stratified by T2DM status and follow-up visit participation. The workflow is divided into three major steps; 1) identifying proteins associated with prevalent or incident T2DM using logistic regression analysis, 2) identifying a panel of proteins for multivariate prediction of incident T2DM using a LASSO bootstrap analysis, followed by internal validation using a separate part of the AGES cohort, and 3) combining genetic data from AGES and summary statistics from the DIAMANTE T2DM GWAS to screen all T2DM-associated proteins for potential causality using a Mendelian randomization analysis. pT2DM, prevalent T2DM; iT2DM, incident T2DM in participants with AGESII follow-up visit; iT2DMrec, incident T2DM in participants without AGESII follow-up visit.

**Fig. S2.**
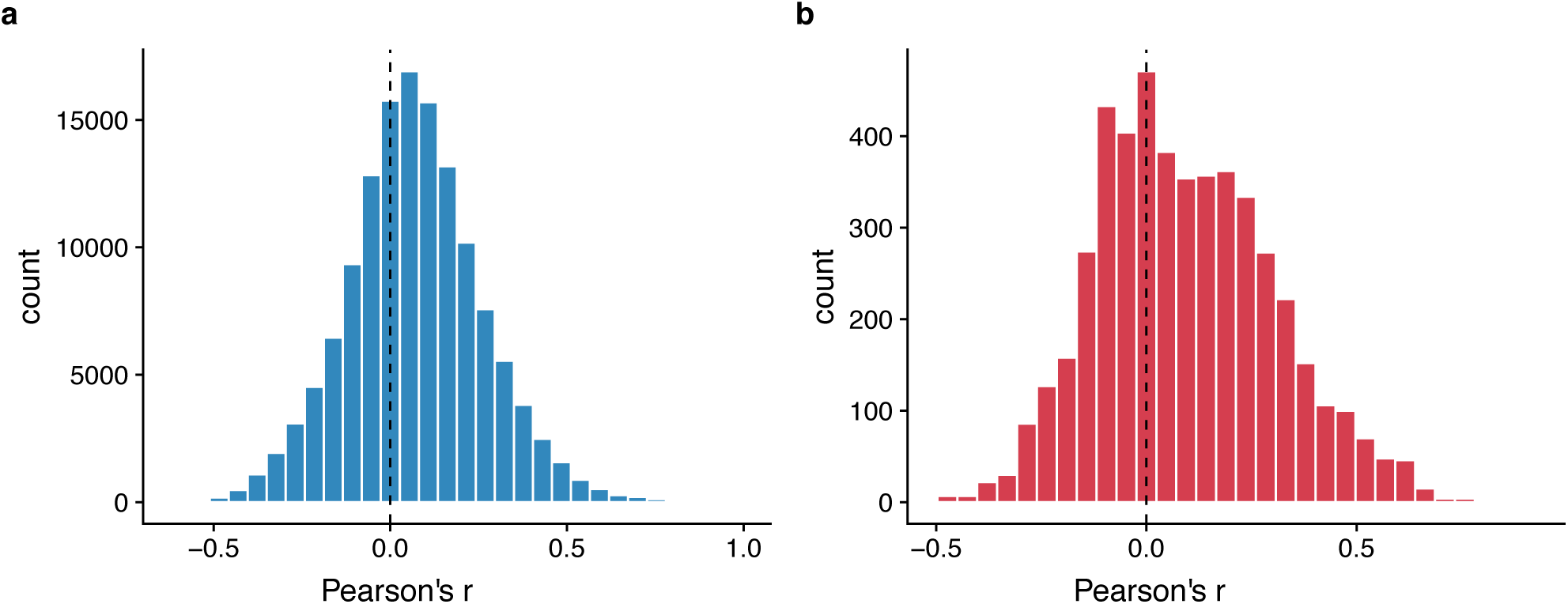
Distribution of Pearson’s correlation coefficients (r) for pairwise correlations between proteins significantly associated with **a)** prevalent T2DM and **b)** incident T2DM.

**Fig. S3.**
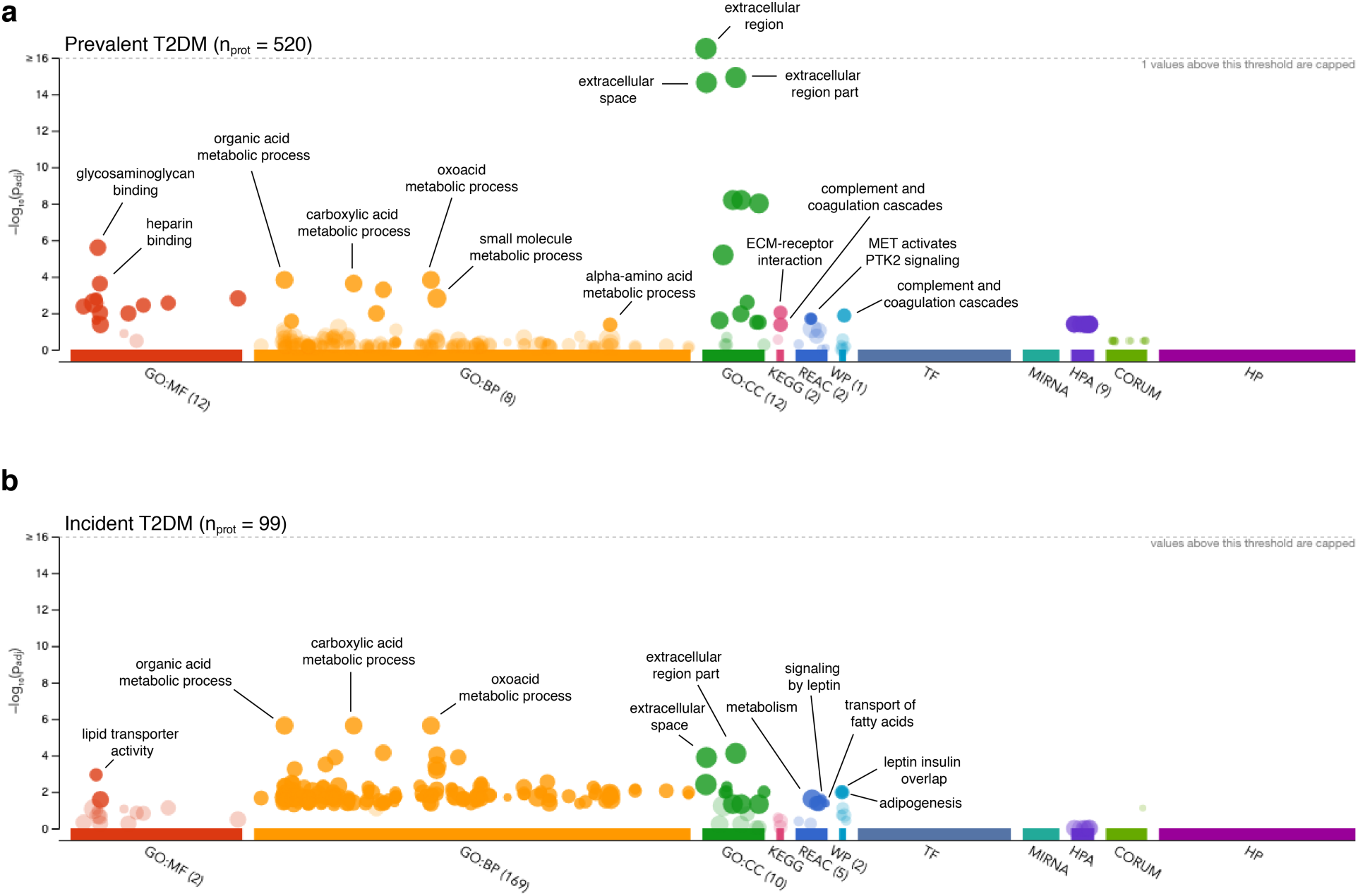
Functional enrichment results from gProfiler for **a)** 520 proteins associated with prevalent T2DM and **b)** 99 proteins associated with incident T2DM.

**Fig. S4.**
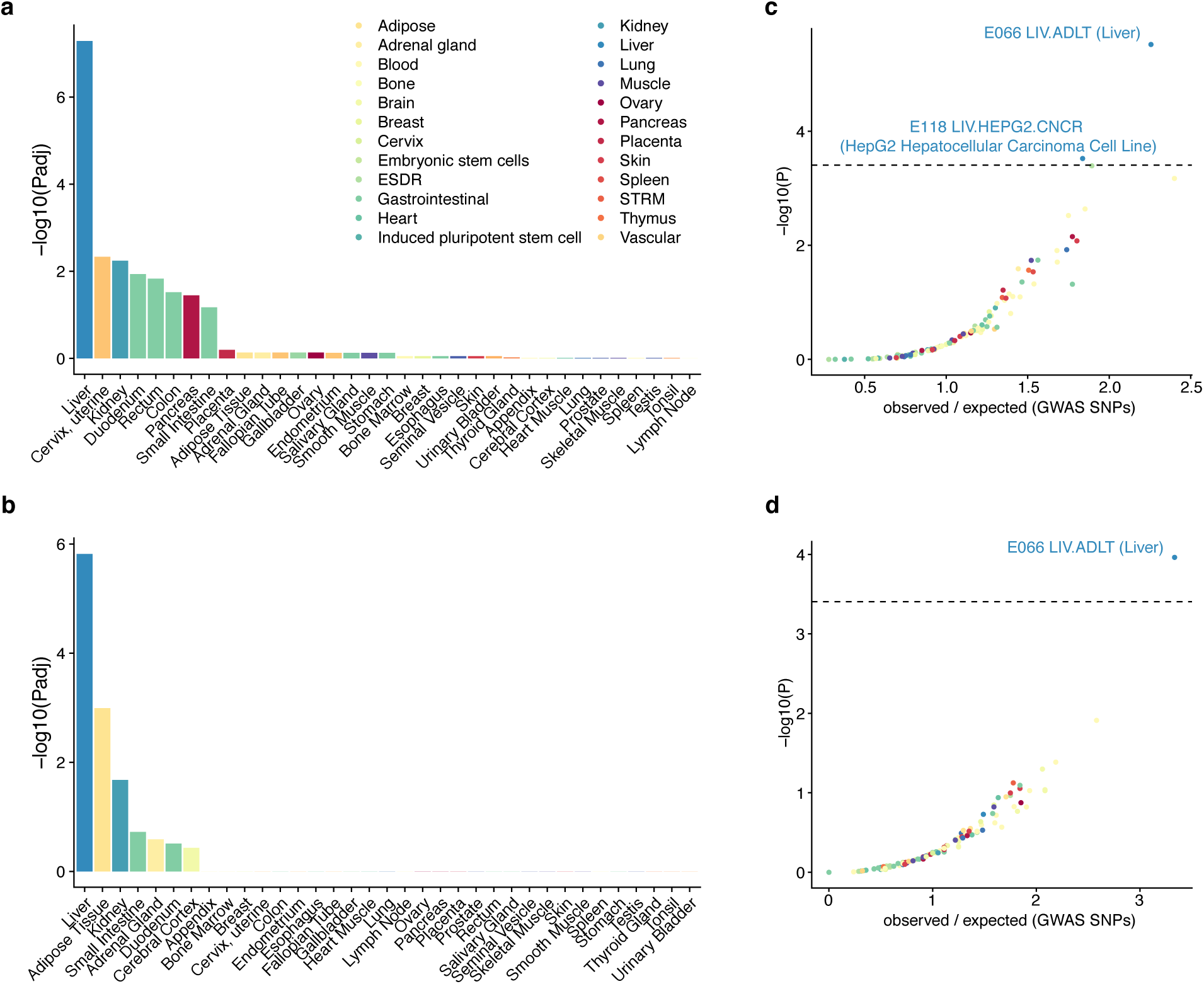
**a)** Tissue-specific gene expression enrichment for the 520 proteins associated with prevalent T2DM compared to the full panel of 4,137 proteins measured, **b)** Tissue-specific gene expression enrichment for 99 proteins associated with incident T2DM compared to the full panel of 4,137 proteins measured, **c)** Cell-type specific enhancer enrichment of genetic variants regulating levels of proteins associated with prevalent T2DM compared to GWAS SNPs, **d)** Cell-type specific enhancer element enrichment of genetic variants regulating levels of proteins associated with incident T2DM compared to GWAS SNPs.

**Fig. S5.**
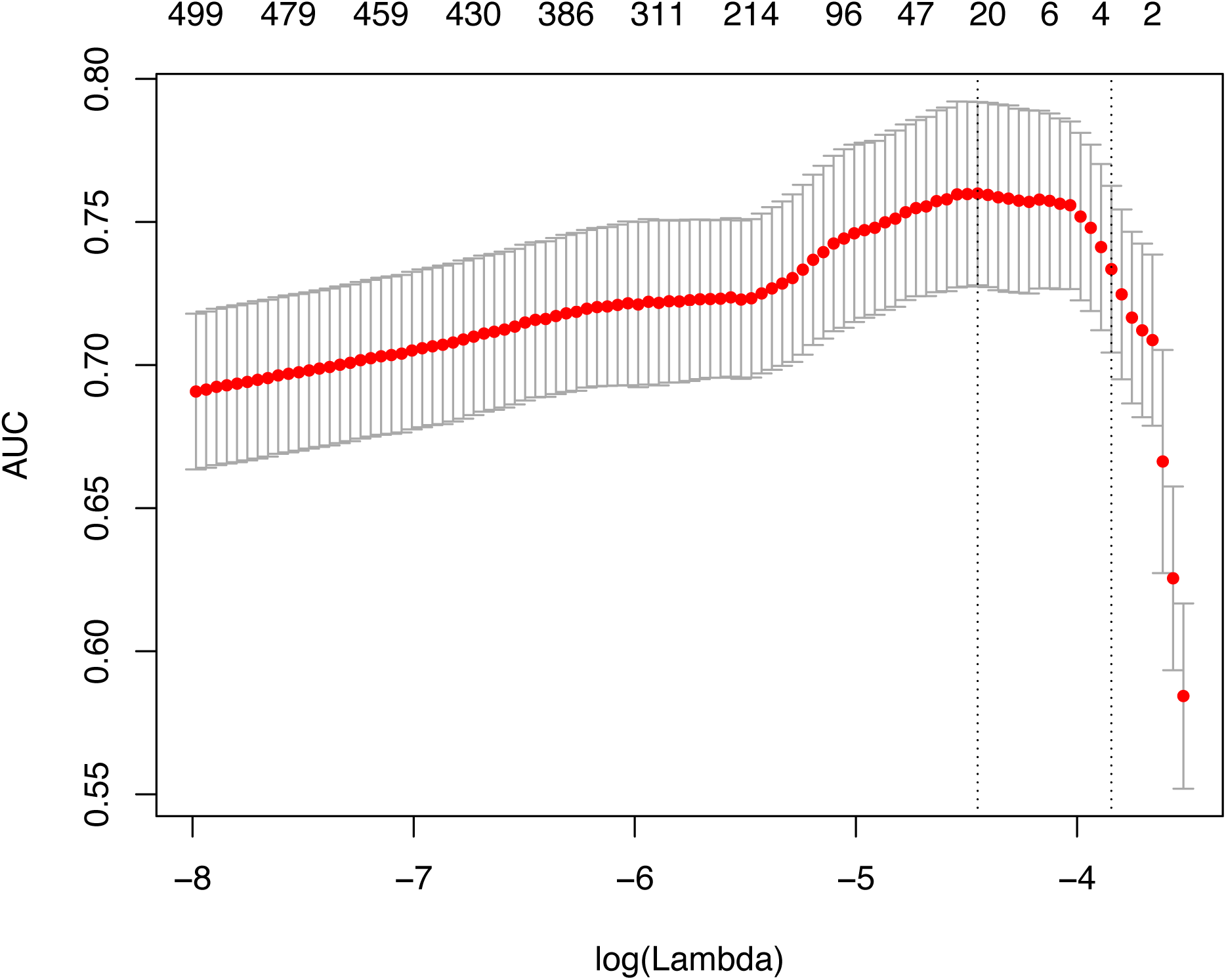
An example of a LASSO regression output for incident T2DM (n = 2,940, n_case_ = 112). A set of 27 non-zero parameter estimates gave the highest AUC when the tuning parameter log(lambda) was −4.54.

**Fig. S6.**
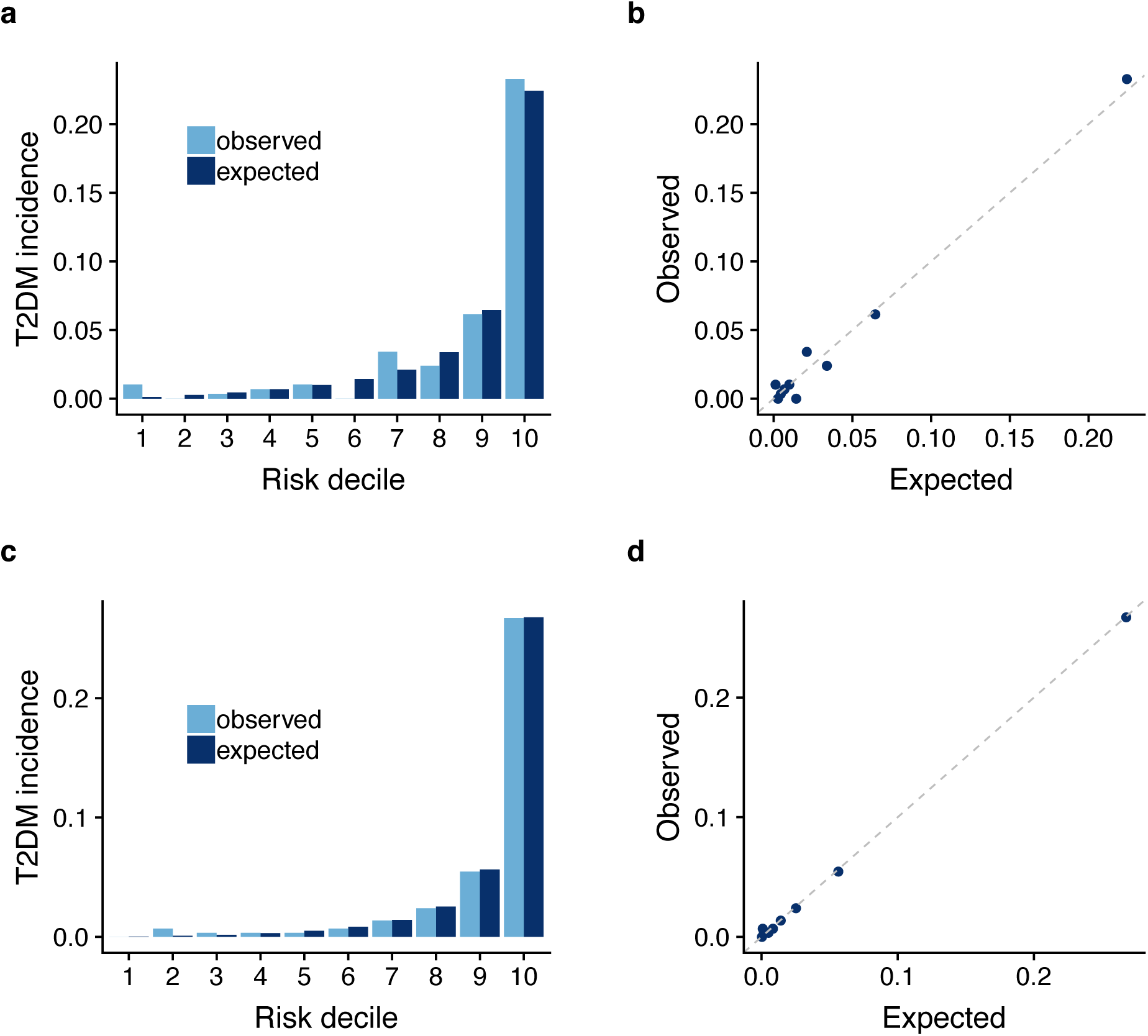
Calibration plots in the AGES sample with 5-year follow-up data (n = 2,940, n_FORS_ = 2,926), showing observed and predicted proportion of individuals with incident T2DM in each risk decile of the discrimination model including **a-b)** the FORS clinical model variables and **c-d)** the FORS clinical model variables plus the top 20 proteins from the LASSO analysis.

**Fig. S7.**
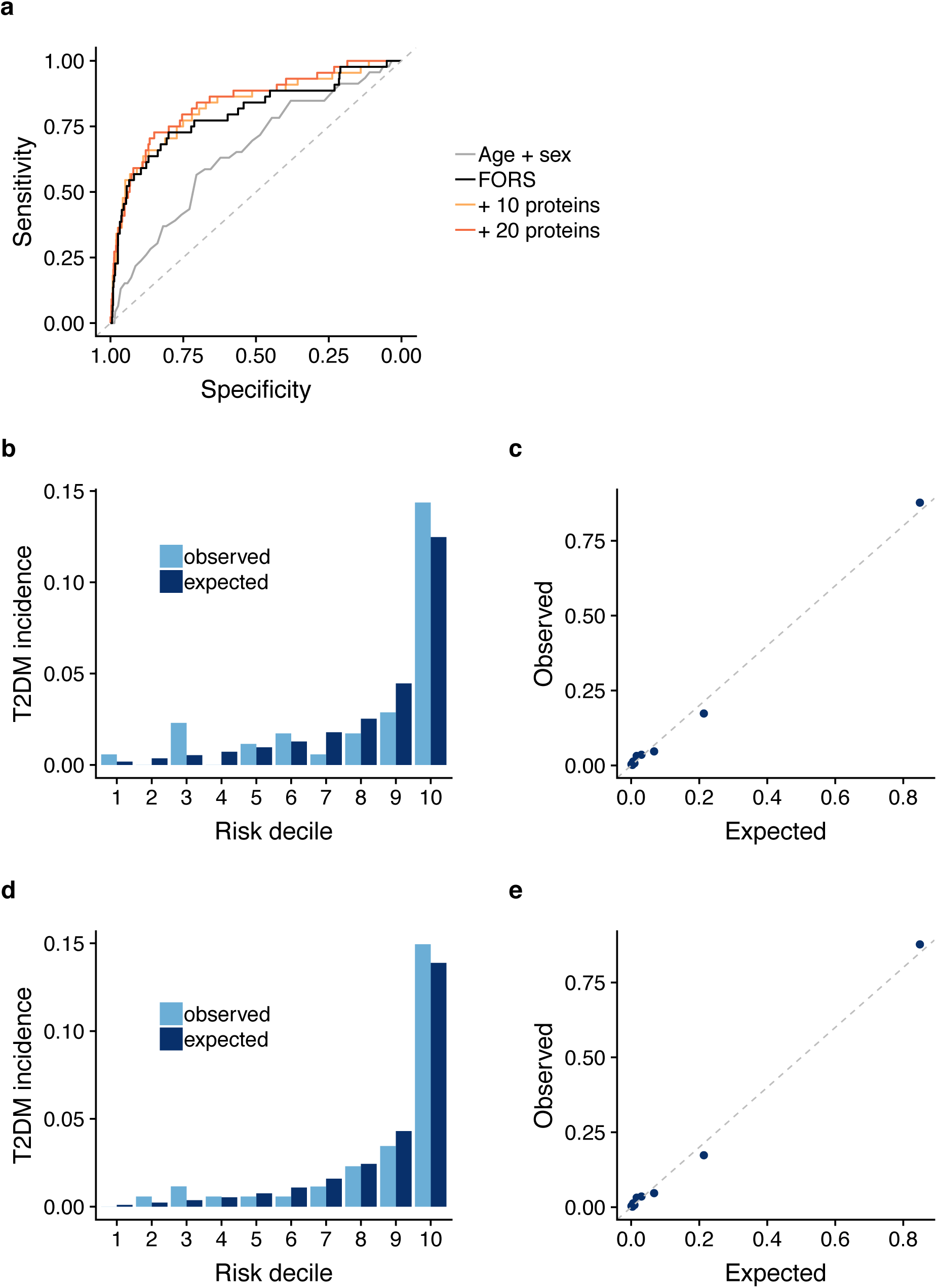
**a)** ROC curves showing the added value of top 10 and 20 ranked proteins (orange shades) for prediction of incident T2DM compared to age and sex (grey) and the Framingham-Offspring risk score, FORS (black) in the AGES validation sample (n = 1,844, n_FORS_ = 1,743). **b-c)** Calibration plots showing observed and predicted proportion of individuals with incident T2DM in the AGES validation sample (n = 1,844) in each risk decile of the discrimination model including the FORS clinical variables and **d-e)** the FORS clinical variables plus the 20 proteins.

**Fig. S8.**
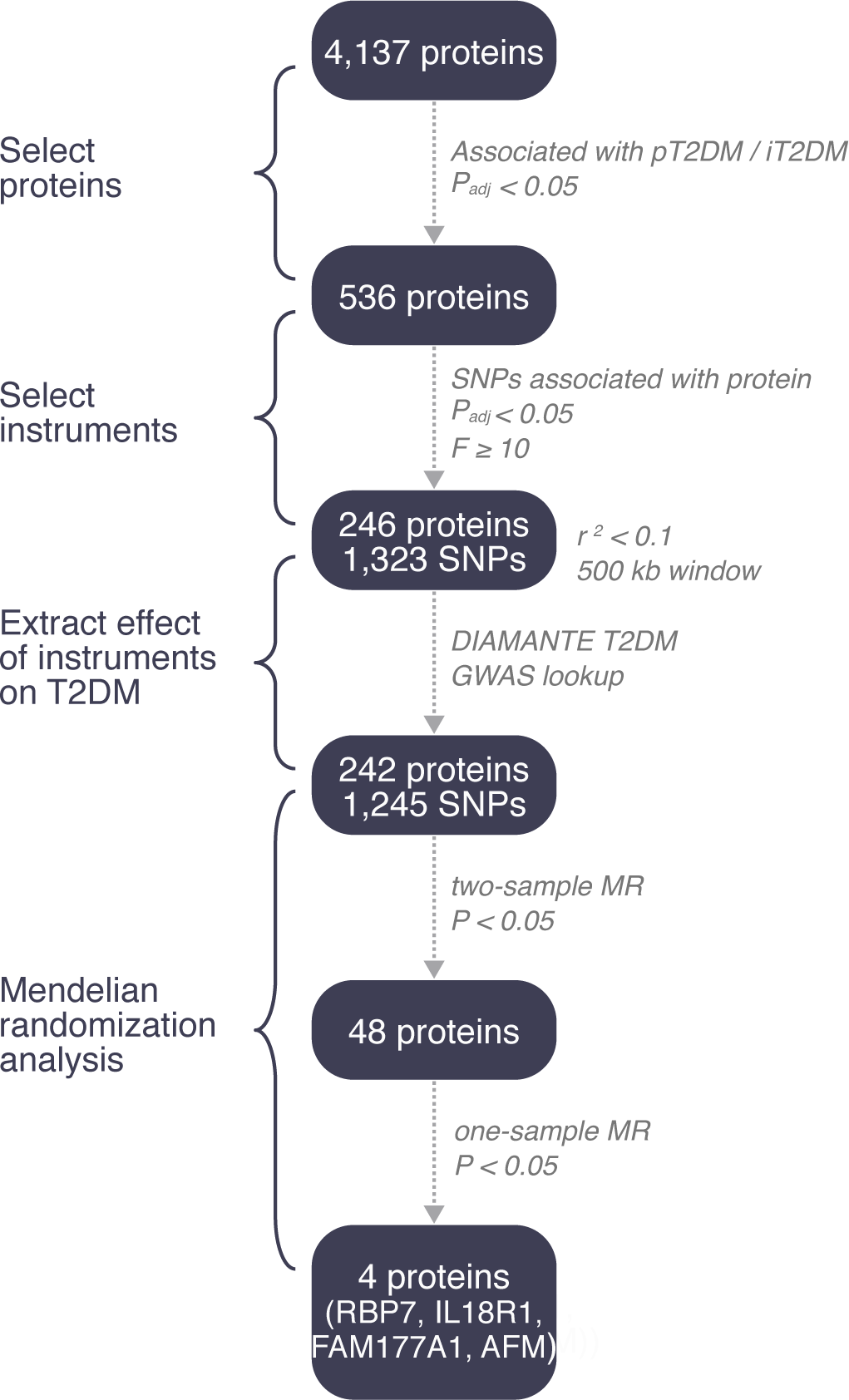
Flowchart illustrating the main steps of the Mendelian randomization analysis for proteins associated with incident or prevalent T2DM in the AGES cohort.

**Fig. S9.**
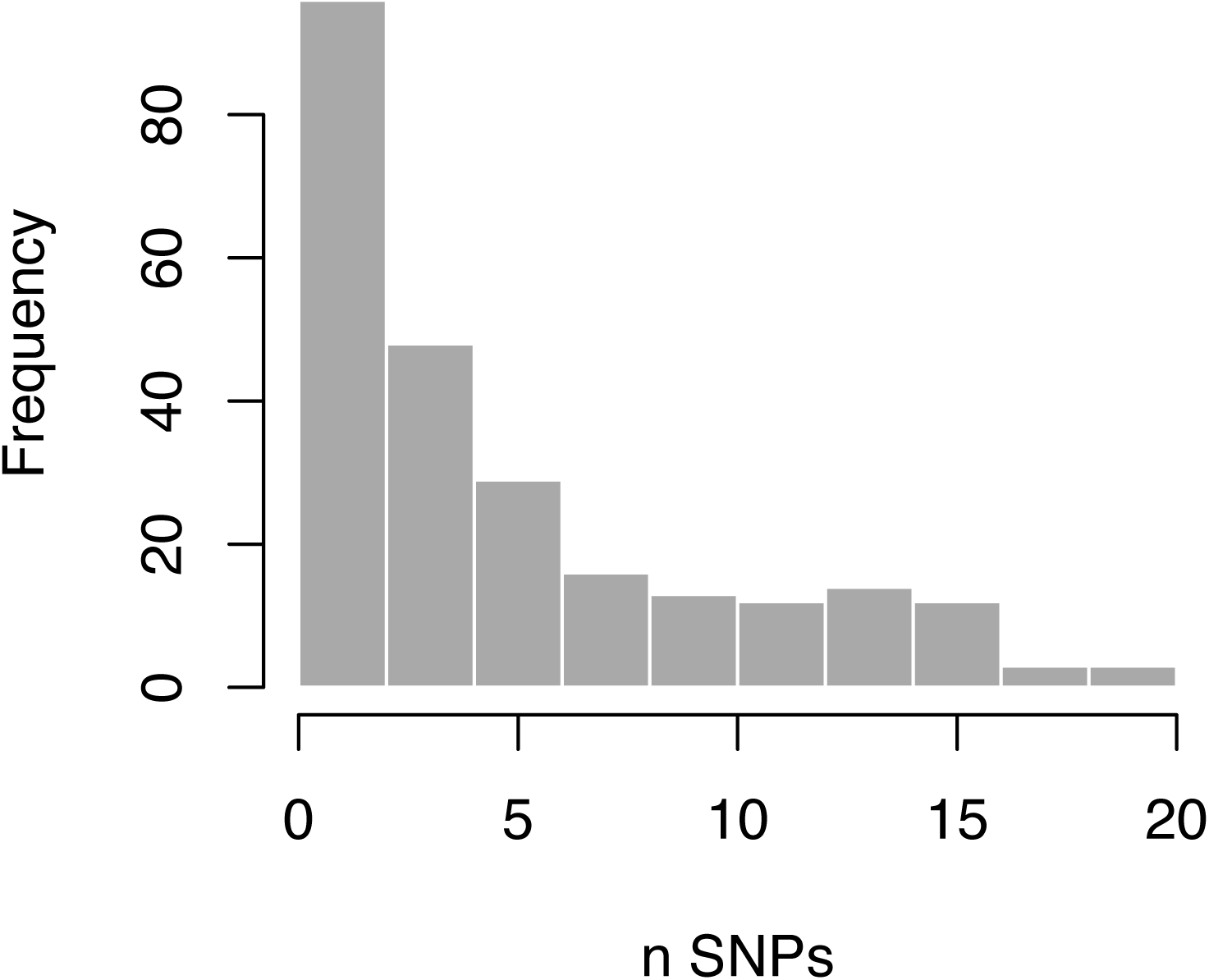
Histogram for the number of instruments identified per protein in the AGES cohort. Independent instruments were defined as genetic variants not in LD (r^2^ < 0.1) and >500 kb apart.

**Fig. S10.**
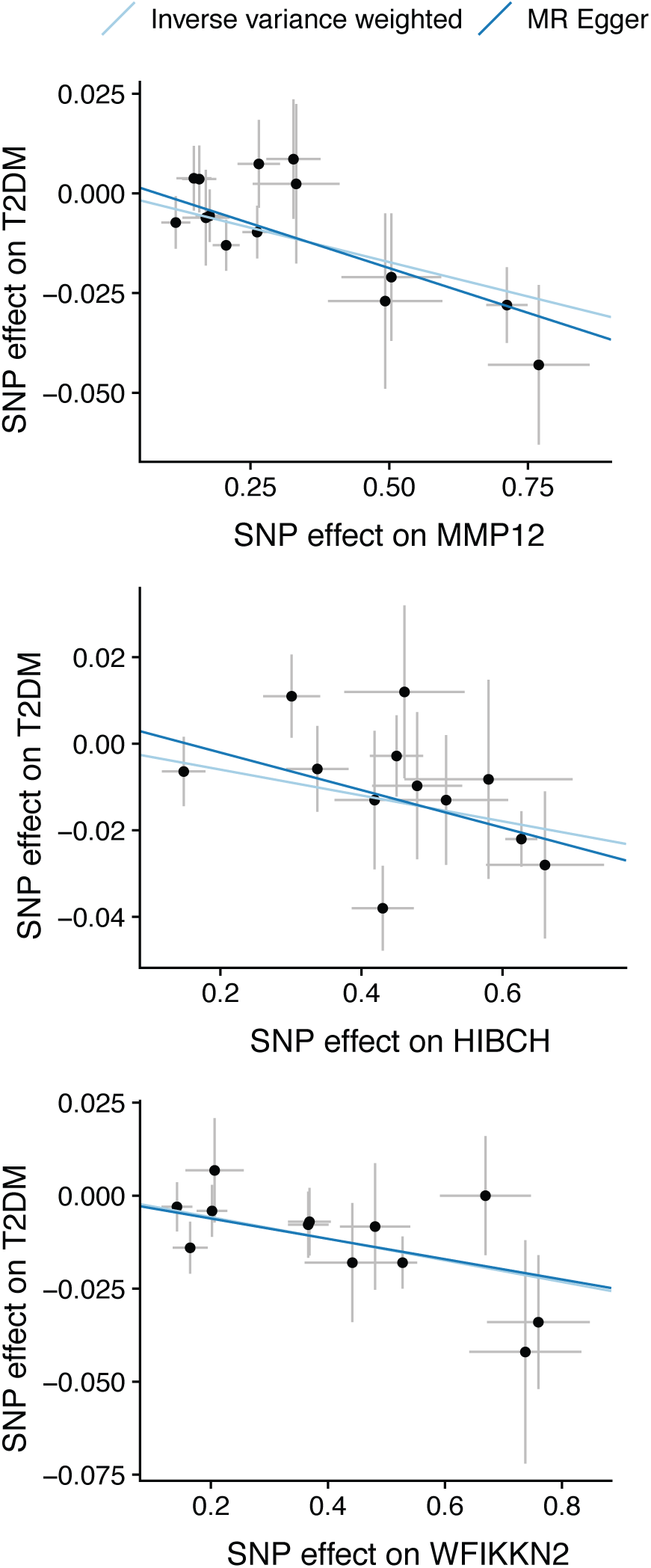
Scatterplots for the top three significant proteins in the two-sample MR, demonstrating the estimated effects (with 95% confidence intervals) of their respective genetic instruments on the protein levels in AGES (x-axis) and the risk of T2DM in the DIAMANTE GWAS (y-axis)

**Fig. S11.**
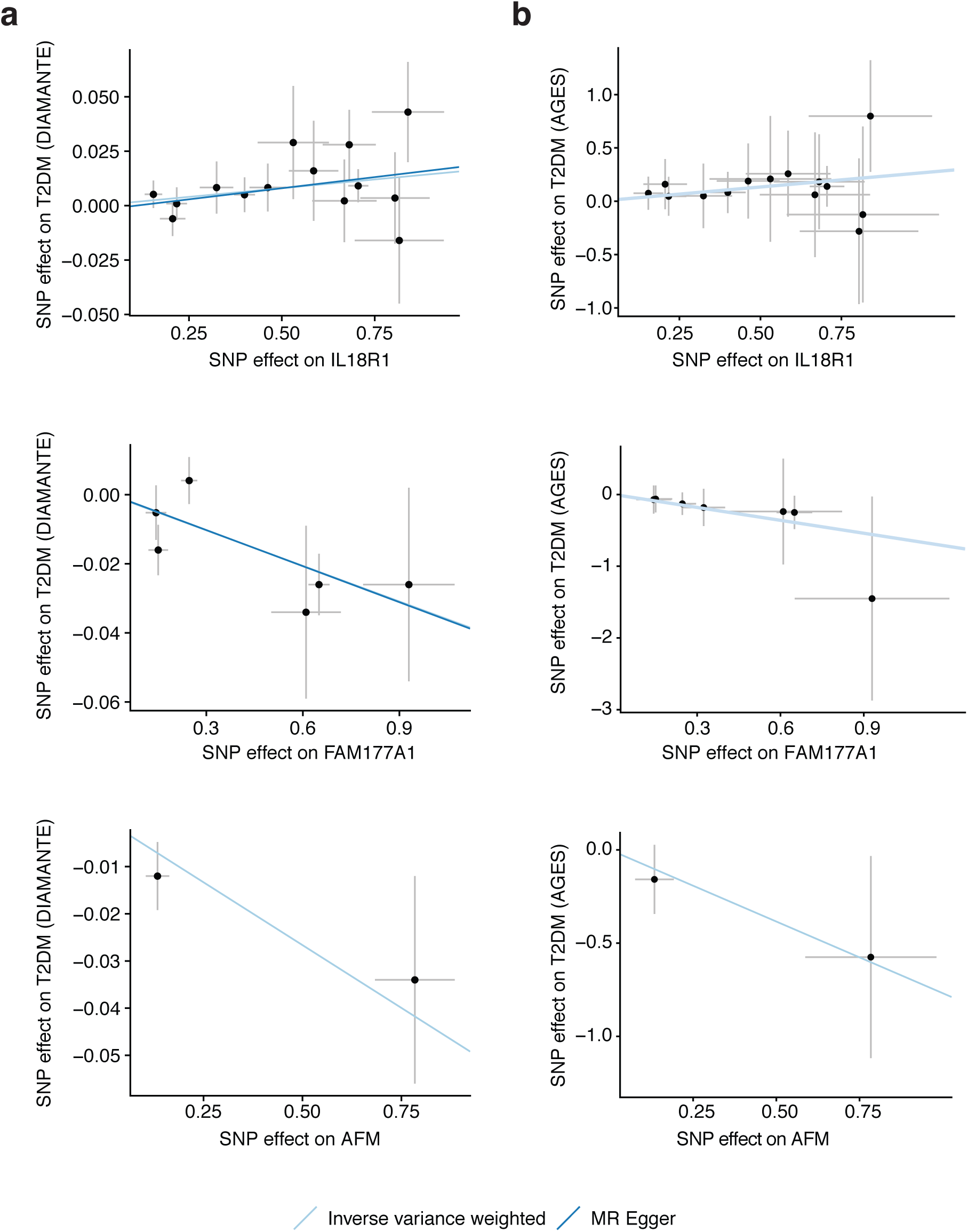
Scatterplots for the three proteins with P < 0.05 and directionally consistent in both the two- and one- sample MR analyses and more than one genetic instrument, demonstrating the estimated effects (with 95% confidence intervals) of their respective genetic instruments on the protein levels in AGES (x-axis) and the risk of T2DM in **a)** the DIAMANTE GWAS (y-axis) and **b)** in the AGES cohort.

**Fig. S12.**
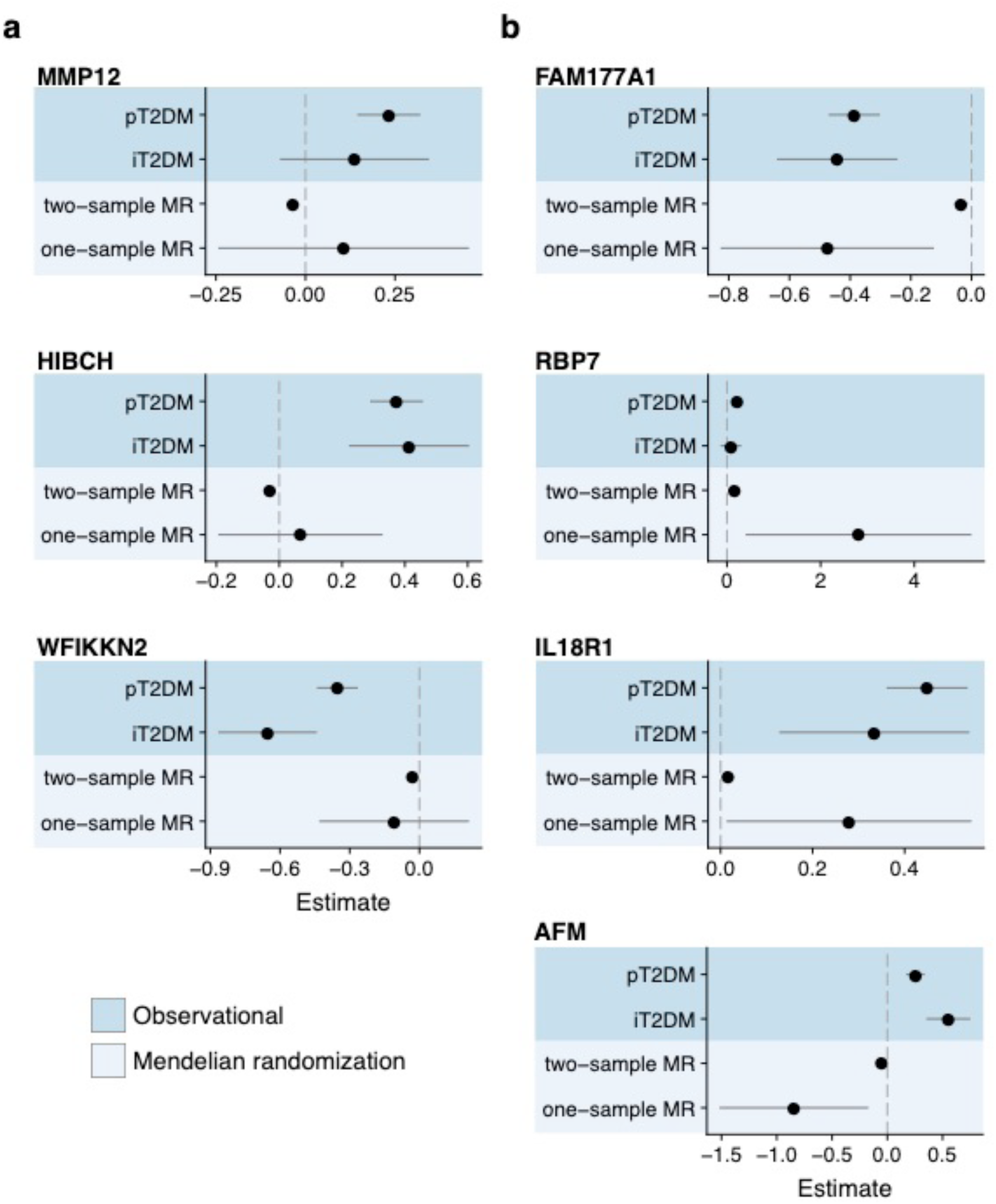
Forest plots comparing observational estimates (darker blue) for incident and prevalent T2DM, and MR estimates (lighter blue) for T2DM in two- and one-sample MR analyses for **a)** the top three significant proteins in the two-sample MR analysis and **b)** the four proteins with P < 0.05 and directionally consistent in both two- and one-sample MR analyses. Error bars represent 95% confidence intervals.

**Fig. S13.**
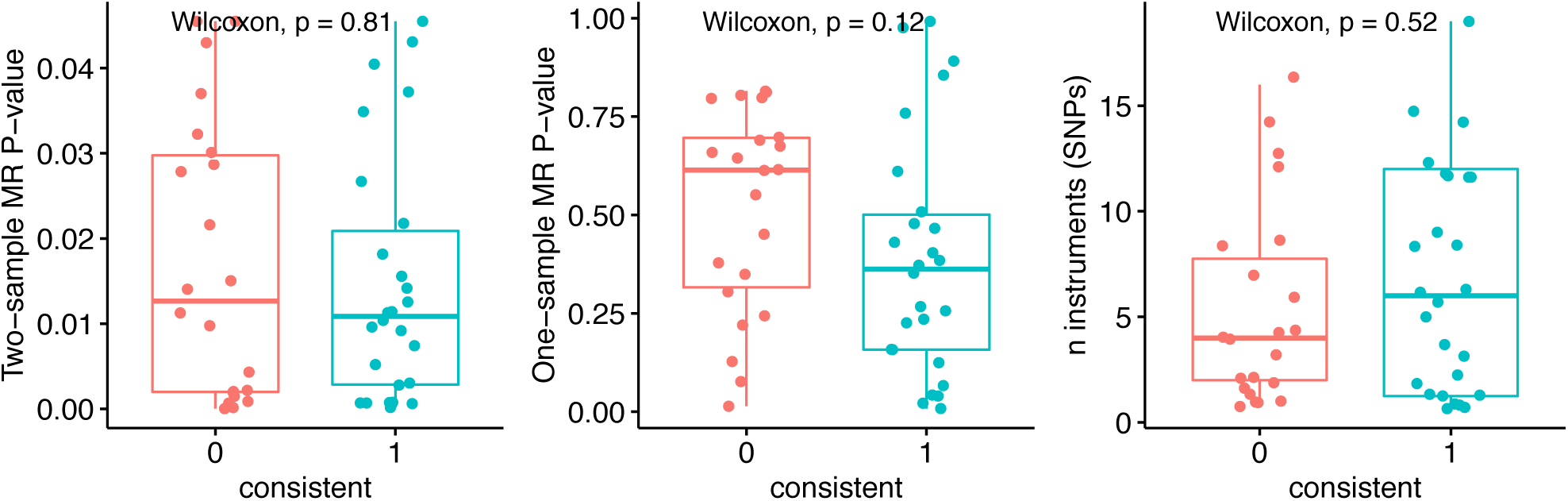
Comparison of MR P-values (two- and one-sample) and number of instruments by directional consistency between observational estimate for prevalent T2DM and two-sample MR.

## References

1. Cho, N. H. et al. IDF Diabetes Atlas: Global estimates of diabetes prevalence for 2017 and projections for 2045. Diabetes Res. Clin. Pract. 138, 271–281 (2018).

2. Morris, A. P. et al. Large-scale association analysis provides insights into the genetic architecture and pathophysiology of type 2 diabetes. Nat. Genet. 44, 981–990 (2012).

3. Mahajan, A. et al. Genome-wide trans-ancestry meta-analysis provides insight into the genetic architecture of type 2 diabetes susceptibility. Nat. Genet. 46, 234–44 (2014).

4. Fuchsberger, C. et al. The genetic architecture of type 2 diabetes. Nature 536, 41–7 (2016).

5. Scott, R. A. et al. An Expanded Genome-Wide Association Study of Type 2 Diabetes in Europeans. Diabetes 147–148 (2017). doi:10.2337/db16-1253

6. Mahajan, A. et al. Fine-mapping type 2 diabetes loci to single-variant resolution using high-density imputation and islet-specific epigenome maps. Nat. Genet. 50, 1505–1513 (2018).

7. Khera, A. V. et al. Genome-wide polygenic scores for common diseases identify individuals with risk equivalent to monogenic mutations. Nat. Genet. (2018). doi:10.1038/s41588-018-0183-z

8. Roberts, L. D., Koulman, A. & Griffin, J. L. Towards metabolic biomarkers of insulin resistance and type 2 diabetes: Progress from the metabolome. Lancet Diabetes Endocrinol. 2, 65–75 (2014).

9. Merino, J. et al. Metabolomics insights into early type 2 diabetes pathogenesis and detection in individuals with normal fasting glucose. Diabetologia 61, 1315–1324 (2018).

10. Abbasi, A. et al. A systematic review of biomarkers and risk of incident type 2 diabetes: An overview of epidemiological, prediction and aetiological research literature. PLoS One 11, (2016).

11. Davies, D. R. et al. Unique motifs and hydrophobic interactions shape the binding of modified DNA ligands to protein targets. Proc. Natl. Acad. Sci. 109, 19971–19976 (2012).

12. Hathout, Y. et al. Large-scale serum protein biomarker discovery in Duchenne muscular dystrophy. Proc. Natl. Acad. Sci. 112, 7153–7158 (2015).

13. Gold, L. et al. Aptamer-Based Multiplexed Proteomic Technology for Biomarker Discovery. PLoS One 5, e15004 (2010).

14. Emilsson, V. et al. Coregulatory networks of human serum proteins link genetics to disease. Science (80-.). 1327, 1–12 (2018).

15. Harris, T. B. et al. Age, gene/environment susceptibility-Reykjavik study: Multidisciplinary applied phenomics. Am. J. Epidemiol. 165, 1076–1087 (2007).

16. Wilson, P. W. F. et al. Prediction of incident diabetes mellitus in middle-aged adults: The Framingham Offspring Study. Arch. Intern. Med. 167, 1068–1074 (2007).

17. Smith, G. D. & Hemani, G. Mendelian randomization: Genetic anchors for causal inference in epidemiological studies. Hum. Mol. Genet. 23, 89–98 (2014).

18. Goncalves, I. et al. Elevated plasma levels of MMP-12 are associated with atherosclerotic burden and symptomatic cardiovascular disease in subjects with type 2 diabetes. Arterioscler. Thromb. Vasc. Biol. 35, 1723–1731 (2015).

19. Sun, B. B. et al. Genomic atlas of the human plasma proteome. Nature 558, 73–79 (2018).

20. Mahdessian, H. et al. Integrative studies implicate matrix metalloproteinase-12 as a culprit gene for large-artery atherosclerotic stroke. J. Intern. Med. 282, 429–444 (2017).

21. Nowak, C. et al. Protein biomarkers for insulin resistance and type 2 diabetes risk in two large community cohorts. Diabetes 65, 276–284 (2016).

22. Raynor, L. A. et al. Novel risk factors and the prediction of type 2 diabetes in the Atherosclerosis Risk in Communities (ARIC) study. Diabetes Care 36, 70–76 (2013).

23. Herder, C. et al. Immunological and cardiometabolic risk factors in the prediction of type 2 diabetes and coronary events: MONICA/KORA Augsburg case-cohort study. PLoS One 6, (2011).

24. Chao, C. et al. The Lack of Utility of Circulating Biomarkers of Inflammation and Endothelial Dysfunction for Type 2 Diabetes Risk Prediction Among Postmenopausal Women. Arch. Intern. Med. 170, 1557–65 (2010).

25. Salomaa, V. et al. Thirty-one novel biomarkers as predictors for clinically incident diabetes. PLoS One 5, 1–8 (2010).

26. Law, B., Fowlkes, V., Goldsmith, J. G., Carver, W. & Goldsmith, E. C. Diabetesinduced alterations in the extracellular matrix and their impact on myocardial function. Microsc. Microanal. 18, 22–34 (2012).

27. Guasch-Ferré, M. et al. Metabolomics in prediabetes and diabetes: A systematic review and meta-analysis. Diabetes Care 39, 833–846 (2016).

28. White, P. J. & Newgard, C. B. Branched-chain amino acids in disease. Science (80-.). 363, 582–583 (2019).

29. Furuhashi, M. & Hotamisligil, G. S. Fatty acid-binding proteins: Role in metabolic diseases and potential as drug targets. Nat. Rev. Drug Discov. 7, 489–503 (2008).

30. Rosell, M. et al. Peroxisome proliferator-activated receptors-α and -γ and cAMPmediated pathways, control retinol-binding protein-4 gene expression in brown adipose tissue. Endocrinology 153, 1162–1173 (2012).

31. Tamori, Y., Sakaue, H. & Kasuga, M. RBP4, an unexpected adipokine. Nat. Med. 12, 30–31 (2006).

32. Hsiao, G. et al. Multi-tissue, selective PPAR modulation of insulin sensitivity and metabolic pathways in obese rats. AJP Endocrinol. Metab. 300, E164–E174 (2011).

33. Kollerits, B. et al. Plasma Concentrations of Afamin Are Associated With Prevalent and Incident Type 2 Diabetes: A Pooled Analysis in More Than 20,000 Individuals. Diabetes Care 40, 1386–1393 (2017).

34. Kondás, K., Szláma, G., Trexler, M. & Patthy, L. Both WFIKKN1 and WFIKKN2 have high affinity for growth and differentiation factors 8 and 11. J. Biol. Chem. 283, 23677–84 (2008).

35. Li, H. et al. GDF11 attenuates development of type 2 diabetes via improvement of islet β-cells function and survival. Diabetes 66, 1914–1927 (2017).

36. Guo, T. et al. Myostatin inhibition prevents diabetes and hyperphagia in a mouse model of lipodystrophy. Diabetes 61, 2414–23 (2012).

37. American Diabetes Association. Diagnosis and classification of diabetes mellitus. Diabetes Care 37, S81–90 (2014).

38. Candia, J. et al. Assessment of Variability in the SOMAscan Assay. Sci. Rep. 7, 14248 (2017).

39. Psaty, B. M. et al. Cohorts for Heart and Aging Research in Genomic Epidemiology (CHARGE) Consortium. Circ. Cardiovasc. Genet. 2, 73–80 (2009).

40. Kuhn, M. & Johnson, K. Applied Predictive Modeling. (2013). doi:10.1007/978-1-4614-6849-3

41. Reimand, J. et al. g:Profiler—a web server for functional interpretation of gene lists (2016 update). Nucleic Acids Res. 44, W83–W89 (2016).

42. Eden, E., Navon, R., Steinfeld, I., Lipson, D. & Yakhini, Z. GOrilla: a tool for discovery and visualization of enriched GO terms in ranked gene lists. BMC Bioinformatics 10, 48 (2009).

43. Jain, A. & Tuteja, G. TissueEnrich: Tissue-specific gene enrichment analysis. Bioinformatics (2018). doi:doi: 10.1093/bioinformatics/bty890

44. Hanley, A. J. G. et al. Prediction of type 2 diabetes using simple measures of insulin resistance: combined results from the San Antonio Heart Study, the Mexico City Diabetes Study, and the Insulin Resistance Atherosclerosis Study. Diabetes 52, 463–9 (2003).

45. Robin, X. et al. pROC: an open-source package for R and S+ to analyze and compare ROC curves. BMC Bioinformatics 12, 77 (2011).

46. Hemani, G. et al. The MR-Base platform supports systematic causal inference across the human phenome. Elife 7, e34408 (2018).

47. Ward, L. D. & Kellis, M. HaploReg v4: Systematic mining of putative causal variants, cell types, regulators and target genes for human complex traits and disease. Nucleic Acids Res. 44, D877–D881 (2016).

